# MIND Networks: Robust Estimation of Structural Similarity from Brain MRI

**DOI:** 10.1101/2022.10.12.511922

**Authors:** Isaac Sebenius, Jakob Seidlitz, Varun Warrier, Richard A I Bethlehem, Aaron Alexander-Bloch, Travis T Mallard, Rafael Romero Garcia, Edward T Bullmore, Sarah E Morgan

**Affiliations:** Department of Psychiatry, University of Cambridge, Cambridge, United Kingdom; Department of Computer Science and Technology, University of Cambridge, Cambridge, United Kingdom; Department of Psychiatry, University of Pennsylvania, Philadelphia, PA, USA; Department of Child and Adolescent Psychiatry and Behavioral Science, The Children’s Hospital of Philadelphia, Philadelphia, PA, USA; Lifespan Brain Institute, The Children’s Hospital of Philadelphia, Philadelphia, PA, USA; Autism Research Centre, Department of Psychiatry, University of Cambridge, Cambridge, United Kingdom; Department of Psychiatry, Harvard Medical School, Boston MA, USA; Psychiatric and Neurodevelopmental Genetics Unit, Center for Genomic Medicine, Massachusetts General Hospital, Boston MA, USA; Instituto de Biomedicina de Sevilla (IBiS) HUVR/CSIC/Universidad de Sevilla/CIBERSAM, ISCIII, Dpto. de Fisiología Médica y Biofísica, Spain; The Alan Turing Institute, London, United Kingdon

**Keywords:** Neuroimaging, T1-weighted MRI, network neuroscience, connectivity, multi-modal, PRIME-IDE, genetics, connectomics, cross-species, macaque, cortical structure

## Abstract

Structural similarity networks are a central focus of magnetic resonance imaging (MRI) research into human brain connectomes in health and disease. We present Morphometric INverse Divergence (MIND), a robust method to estimate within-subject structural similarity between cortical areas based on the Kullback-Leibler divergence between the multivariate distributions of their structural features. Compared to the prior approach of morphometric similarity networks (MSNs) on N>10,000 data from the ABCD cohort, MIND networks were more consistent with known cortical symmetry, cytoarchitecture, and (in N=19 macaques) gold-standard tract-tracing connectivity, and were more invariant to cortical parcellation. Importantly, MIND networks were remarkably coupled with cortical gene co-expression, providing fresh evidence for the unified architecture of brain structure and transcription. Using kinship (N=1282) and genetic data (N=4085), we characterized the heritability of MIND phenotypes, identifying stronger genetic influence on the relationship between structurally divergent regions compared to structurally similar regions. Overall, MIND presents a biologically-validated lens for analyzing the structural organization of the cortex using readily-available MRI measurements.

## Introduction

A single structural MRI scan of a human brain contains an immense amount of information. Standard MRI-based surface reconstructions of the cortex, for example, are comprised of hundreds of thousands of vertices, each characterized by many parameters (Fischl, 2012). The challenging task of integrating this wealth of information to model the structural architecture of the brain is essential to better understand healthy and disordered brain development and function.

Traditional, univariate studies of brain structure focus on changes in individual structural features, such as cortical thickness and volume, with recent large-scale research in this vein revealing the central role of MRI-derived brain structural features to understanding development and disease (Bethlehem et al, 2022). However, brain regions do not function or develop in isolation but instead form an integrated, genetically-coordinated network structure; accurately modelling this architecture is crucial for understanding its putative role across typical and atypical functioning and development (Bullmore and Sporns, 2009; Warrier et al, 2022b; Taquet et al, 2020). Recently, the construction of structural similarity networks has emerged as a promising approach for integrating multiple structural MRI features into biologically-relevant single-subject connectomes (Seidlitz et al, 2018; Li et al, 2017). Morphometric similarity networks (MSNs), the prevailing such method, are constructed by representing each brain region as a vector of several MRI features (for example, the region’s mean cortical thickness) and using the pairwise correlation of these (*Z*-scored) feature vectors as a measure of the structural similarity between brain regions.

While simple in construction, MSNs have demonstrated the promise of structural similarity networks to link macro-scale MRI phenotypes with their neurobiological substrates. For example, MSNs recapitulated known brain organizational principles and cortical cytoarchitectonic classes (von Economo and Koskinas, 1925) more robustly than comparable networks derived from tractography of diffusion-weighted imaging (DWI) data, or from structural covariance network analysis of cortical thickness, in N~300 healthy young adults (Seidlitz et al, 2018). Moreover, MSNs from macaque MRI data were positively correlated with gold-standard axonal connectivity measured by tract-tracing (Seidlitz et al, 2018). Most promisingly, MSNs have provided a useful bridge between brain structure, cortical gene expression, and genetics. For example, by combining cortical transcriptomic data from the Allen Human Brain Atlas (Hawrylycz et al, 2015) with structural MRI from subjects with one of six different chromosomal copy number variation (CNV) disorders, Seidlitz et al (2020) demonstrated that the changes in morphometric similarity induced by each CNV closely resembled the spatial expression patterning of genes from the affected chromosome. Other studies have shown that changes in morphometric similarity in psychotic disorders (Morgan et al, 2019), major depressive disorder (Li et al, 2021), and Alzheimer’s disease (Zhang et al, 2021) correspond to the cortical expression of disease-relevant genes.

Despite the promise of MSNs, one key limitation is that they reduce the rich, vertex-level data from MRI-based cortical surface reconstructions to single summary statistics for each feature per region. While other work has explored structural similarity measured directly from vertex-level data, these methods were limited to the use of a single structural feature such as cortical thickness (Homan et al, 2019) or grey matter volume (Leming et al, 2021; Kong et al, 2015).

Here, we propose Morphometric INverse Divergence (MIND) as a novel method for effectively estimating structural similarity in the brain. We first characterize each brain region as a multidimensional distribution of a number of structural features at the vertex-level (e.g. vertex-wise cortical thickness values). We then calculate the MIND structural similarity metric based on the symmetric Kullback-Leibler (KL) divergence between the multivariate distributions of each region pair.

In order to comparatively evaluate MSNs and MIND networks, we studied how accurately and consistently each type of network models a wide array of known biological patterns of brain organization. In comparison to MSNs, we show that MIND networks more accurately represented known cortical symmetry and cytoarchitecture, and mapped more closely onto axonal connectivity from retrograde tract-tracing in macaques. MIND networks were also highly consistent across cortical parcellations and between individuals, and were more resilient to the inclusion of noisy features. Most importantly, MIND networks showed a step-change increase in agreement with networks of cortical gene co-expression compared to MSNs, providing fresh evidence of the unified structural and transcriptomic organization of the brain. Building on this result, we estimated the patterns of twin-based and SNP-based heritability in MIND networks and MSNs at multiple scales of network resolution, demonstrating that MIND network phenotypes were systematically more heritable than MSNs and exhibited similar heritability to other structural features such as sulcal depth. We observed a stronger genetic influence on MIND similarity between structurally divergent regions than between regions with similar morphology, and that on average, areas of insular and primary sensory cortex were more heritable than the MIND network hubs of the association cortex.

In sum, MIND presents a new, validated lens for understanding the organizing principles of cortical structure from a single T1w image. Code for MIND calculation is publicly available at https://github.com/isebenius/MIND.

## Results

### Morphometric INverse Divergence: MIND estimator of structural similarity

The pipeline for constructing MIND networks is shown in Figures 1 and S1. As input, we used the mesh reconstructions of the cortical surface generated from T1w images by Freesurfer’s `recon-all` command (Fischl et al, 1999). This surface can be described by a set of vertices (163,842 vertices per hemisphere for the fsaverage template (Fischl, 2012)). Each vertex is characterized by multiple metrics derived from the reconstruction of the MRI image, such as cortical thickness (CT) and sulcal depth (SD), among others. We refer to these derived metrics as structural features.

**Fig. 1.**
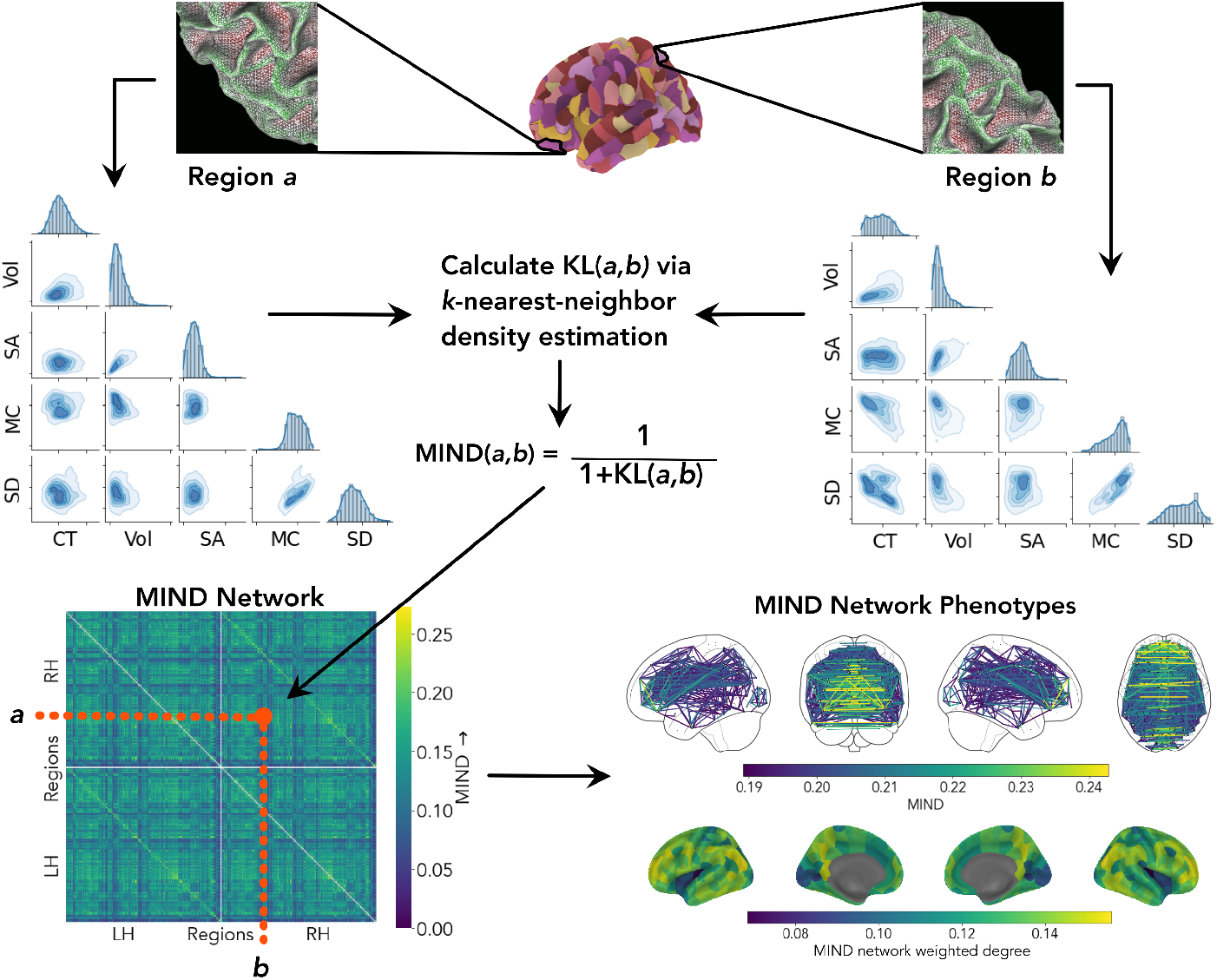
Estimation of Morphometric Inverse Divergence. The cortex is parcellated into a set of predefined regions or areas, each of which comprises a set of vertices themselves described by multiple structural features. Each cortical area is thus characterized by the multidimensional distribution of the structural features measured at each of its constituent vertices. Pairwise KL divergence between these regional multidimensional distributions is estimated using *k*-nearest neighbor density estimation (Perez-Cruz, 2008), and the divergence is then transformed into a similarity score, termed Morphometric Inverse Divergence, 0 < MIND ≤ 1, with higher scores indicating greater similarity. Illustrative distributions for regions *a* and *b* are shown as scatterplot matrices, with diagonal panels showing the marginal univariate distribution for five structural features and the off-diagonals showing each pair-wise bivariate relationship. Bottom row: a visualization of a group-mean MIND similarity matrix, and of the two main MIND network phenotypes (edges and weighted network degrees, calculated as the average edge weight per region) on the brain. The top 2% of MIND edges are shown for visibility. CT: cortical thickness, Vol: grey matter volume, SA: surface area, MC: mean curvature, SD: sulcal depth.

To estimate MIND, we standardize each feature across all vertices, then aggregate the vertices within each region to create a single distribution per region, which is multidimensional due to the inclusion of several structural features. We then compile a pairwise distance matrix by estimating the symmetric Kullback-Leibler (KL) divergence (Kullback and Leibler, 1951) between the multivariate distributions of each pair of cortical regions. Finally, given regions *a* and *b*, we transform their corresponding divergence *KL*(*a*, *b*) as follows to calculate the MIND metric of similarity, bounded between 0 and 1:

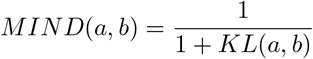

A more rigorous definition of MIND as a similarity metric, in addition to a description of the *k*-nearest neighbor algorithm used to estimate multivariate KL divergence (Perez-Cruz, 2008), is provided in Methods.

### Data and network construction

As our principal human MRI dataset, we used data from 10,367 individuals from the Adolescent Brain Cognitive Development (ABCD) study (Hagler et al, 2019). For each subject, we constructed MSNs and MIND networks using a symmetric subdivision of the Desikan-Killiany atlas (Desikan et al, 2006) into 318 parcels of similar volume (unless otherwise noted), henceforth referred to as DK-318 (Romero-Garcia et al, 2012). We used the five morphometric features indicated in Fig. 1 for both MIND network and MSN construction: cortical thickness (CT), mean curvature (MC), sulcal depth (SD), surface area (SA), and grey matter volume (Vol), which is estimated at the vertex level by combining local measurements of thickness and area. We used these features as they are readily available from standard MRI processing pipelines using T1w images alone (Fischl, 2012); as such, we ensure that the method is applicable to most legacy structural data. However, MIND can be extended to include additional volumetric measurements of interest based on other imaging modalities (e.g. such as fractional anisotropy) using volumetric registration and projection onto the surface mesh; for example, using Freesurfer’s `mri_vol2surf` command.

### Network reliability

To compare the reliability of MIND networks and MSNs as measures of brain network organization, we examined the consistency of edge weights and nodal degrees in both types of structural similarity network, applying the same parcellation template to multiple individual scans to assess between-subject consistency, and applying multiple parcellation templates to the same set of scans to assess parcellation consistency. We also evaluated the effect of including uninformative (noise) features into both types of network construction.

#### Between-subject consistency

The group-level MSN and MIND networks were correlated in terms of both edge weights (*r*=0.48, Fig. 2D) and weighted nodal degrees (*r*=0.38). However, MIND networks were substantially more consistent across subjects (Fig. 2E), measured by pairwise correlation of edges (mean pairwise *r*=0.62 vs. 0.38) and degrees (mean pairwise *r*=0.73 vs. 0.45), suggesting that MIND network construction may lead to less noisy estimates of a common structural architecture.

**Fig. 2.**
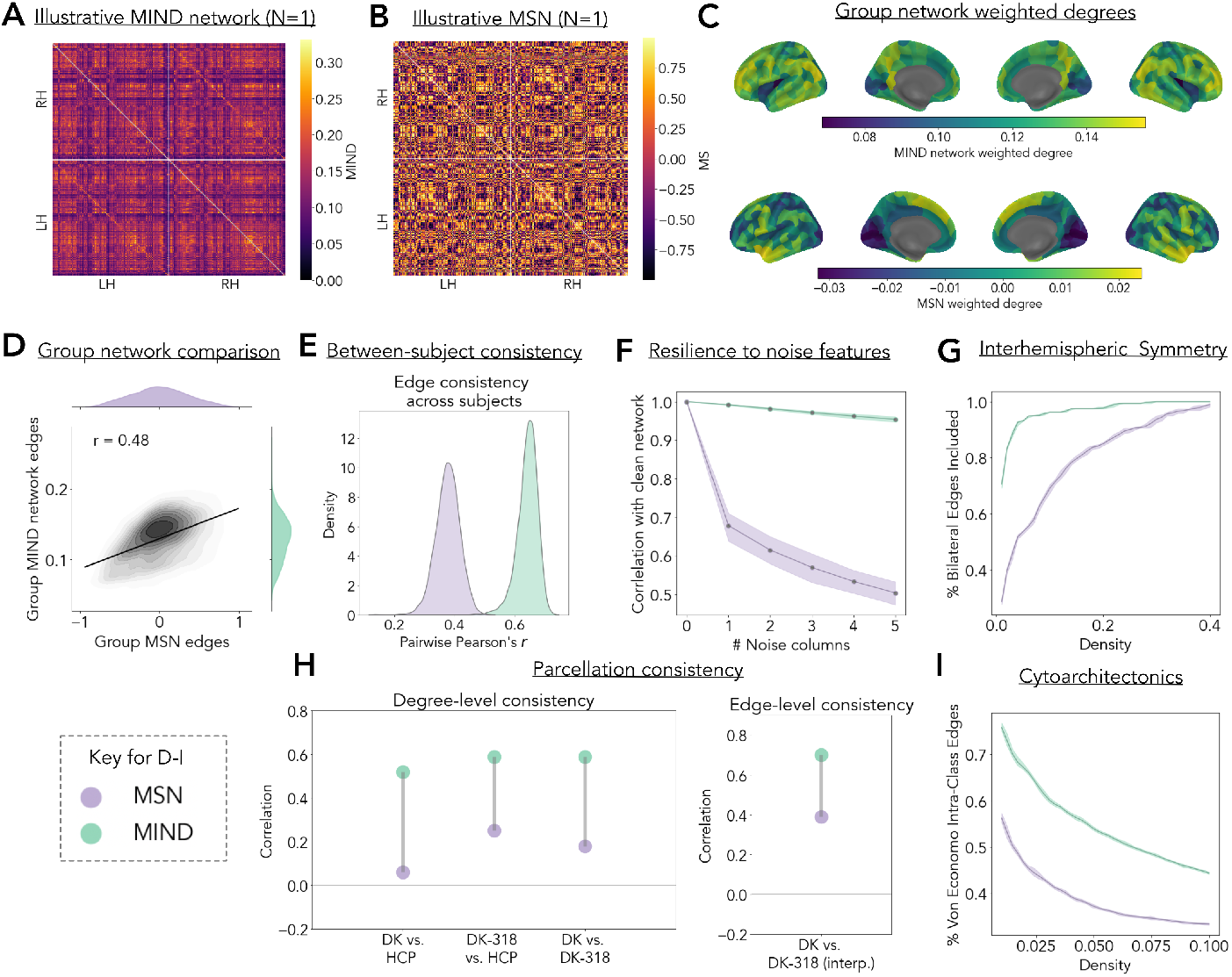
Comparing MIND networks and MSNs. A, B) Illustrative MIND network and MSN from the same random subject in the ABCD cohort. C) Cortical maps of the group-level MIND network and MSN weighted node degree. D) The positive correlation between edge weights of the group level networks. E) The distribution of pairwise correlations of network edges between subjects for MIND networks and MSNs, for all pairs of 10,367 subjects. F) The correlation between MIND networks and MSNs with corresponding networks constructed using 1-5 additional random features of Gaussian noise (for 150 random subjects). Shading represents the empirical 95% CI. G) The fraction of total inter-hemispheric connections represented at different network densities for both types of group mean networks. H) Parcellation consistency of MIND network phenotypes at nodal level (weighted degree) and at edge level. The left plot shows the correlation between weighted degree estimated by one of the possible pairs of parcellation templates: DK, DK-318, and HCP. To calculate between-parcellation correlations, each vertex was assigned the weighted degree of the region within which it fell, for each parcellation, and the correlation was calculated between the resulting vectors of vertex-wise values. The right plot shows the correlation between 2278 network edges calculated using the 68-region Desikan-Killiany (DK) parcellation, or by using the finer-grained DK-318 parcellation to estimate 50,403 edges and coarse-graining (DK-318 interp.) to match the number of edges in the original DK network. I) The fraction of edges between two regional nodes of the same cytoarchitectonic class over a range of network densities. In G) and I), shading represents a 95% CI determined by population bootstrapping. In all panels except as noted in H), the DK-318 parcellation was used to define 318 cortical regions of approximately equal volume.

#### Parcellation consistency

Brain network analysis assumes that major findings can be replicated across cortical parcellations. We analyzed the consistency of group-level MSNs and MIND networks across three commonly used cortical parcellations: the 68-region Desikan-Killiany (DK) atlas (Desikan et al, 2006), the 318-region DK-318 atlas derived by subdivision of DK areas (Romero-Garcia et al, 2012) (the principal parcellation used for this study), and the 360-region HCP parcellation (Glasser et al, 2016).

We examined edge-level consistency by leveraging the fact that DK-318 is a strict subdivision of the DK atlas, allowing us to compare the original group DK networks with interpolated versions derived from the DK-318 group networks (see Methods). MIND networks showed markedly higher edge consistency (Fig. 2H) in terms of the correlation between the original and interpolated DK networks (*r*=0.70 vs. 0.39 for MSNs).

To calculate between-parcellation correlations, each vertex was labeled by the weighted degree of the region to which it was assigned, for each parcellation, and the correlation was estimated between these two identical-length vectors of parcellation-specific degree projected to each vertex (Fig. 2H and Fig. S4). MIND networks were strongly correlated across all (three) possible pairs of the three parcellations, whereas MSN degree demonstrated limited generalizability across parcellations. (e.g. *r*=0.59 vs. 0.18 for MIND networks and MSNs, respectively, when comparing weighted degree for DK and DK-318 atlases).

#### Resilience to noisy features

We studied the robustness of MIND networks and MSNs to the inclusion of uninformative (noise) features. We created additional MIND networks and MSNs with between one and five 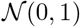 noise features at each vertex (in addition to the five measured MRI features) for a random subset of 150 subjects.^1^ MIND networks constructed from these noisy data were almost perfectly correlated with MIND networks constructed from the measured features only, without additional noise (Figure 2F); whereas MSN construction was significantly degraded by the inclusion of noise features (e.g., mean *r* = 0.95 vs. 0.50 for MIND networks and MSNs with five noise features).

### Validation by principles of cortical organization

We studied the extent to which each network type represented foundational principles known to govern cortical organization. Specifically, we benchmarked the biological validity of each type of structural similarity using the following basic premises about three known principles of brain structure:

- *Symmetry*: the cortex is highly symmetric and homologous regions of right and left hemispheres are reciprocally interconnected, so a valid measure of structural similarity should have strong weights for inter-hemispheric edges.
- *Cortical microstructure*: cortical areas can be cytoarchitectonically classified based on microstructural properties measured histologically, so a valid MRI measure of structural similarity should have strong weights for edges between cortical areas histologically assigned to the same cytoarchitectonic class (von Economo and Koskinas, 1925).
- *Axonal connectivity*: cortical areas are inter-connected by white matter tracts, and cytoarchitectonically similar regions are more likely to be axonally inter-connected, so a valid measure of structural similarity should correlate with axonal connectivity as measured by gold-standard tract-tracing in non-human primates.

#### Symmetry and inter-hemispheric connections

Across a range of network densities, we measured how many bilateral connections were represented by each type of group mean network. Over all densities, MIND networks comprised a substantially larger fraction of bilaterally symmetric connections than MSNs (Fig. 2G).

#### Cytoarchitectonics and within-class connections

Next, we analyzed the extent to which MIND networks and MSNs recapitulated known patterns of cortical microstructure, as measured by higher structural similarity between regions belonging to the same von Economo cytoarchitectonic class (von Economo and Koskinas, 1925). MIND networks demonstrated higher connectivity between regions of the same cytoarchitectonic class across a range of network densities (Fig. 2I), indicating a closer correspondence with known patterns of cytoarchitectonic similarity at the scale of neuronal organization.

#### Axonal connectivity and structural similarity

Previous work has shown that regions with similar cytoarchitecture are more likely to be connected by axonal tracts than regions that are micro-structurally dissimilar (Barbas, 2015; Goulas et al, 2016). We anticipated that more robust estimation of structural similarity via MIND networks, compared to MSNs, would result in stronger correlations with axonal connectivity measured by retrograde tract tracing in the macaque monkey brain.

Using MRI data from 19 macaques (Xu et al, 2020; Milham et al, 2018), we constructed group-level MSN and MIND networks using the same five morphometric features as for human MRI analysis plus the T1/T2 ratio as an estimate of intra-cortical myelination. We compared the correspondence between axonal connectivity and structural similarity across five tract-tracing connectomes (Fig. 3A, see Methods for details).

**Fig. 3.**
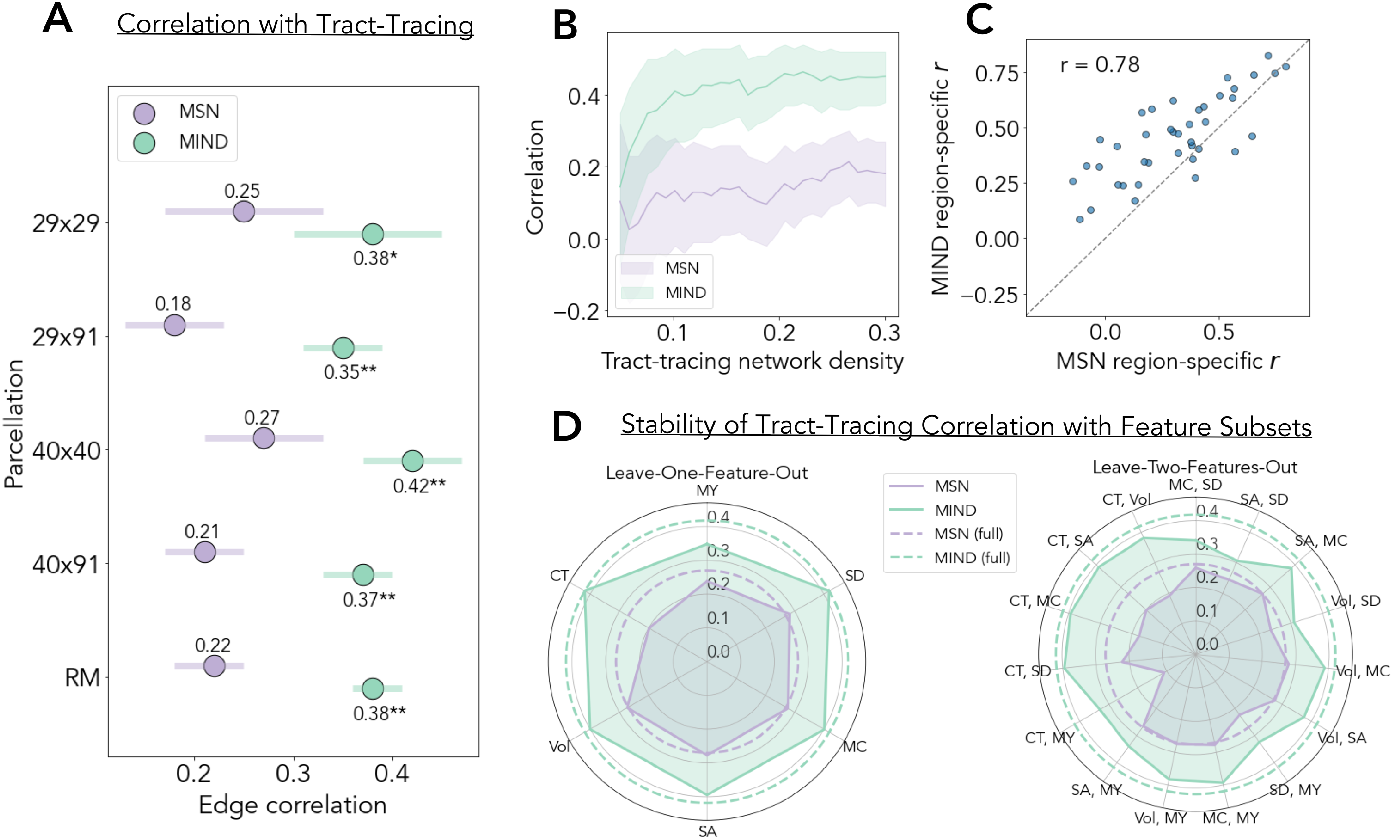
Structural similarity from MRI compared to axonal connectivity from tract-tracing in the macaque brain. A) Correlation between structural similarity edge weights, in MIND networks or MSNs derived from MRI, and axonal connectivity edge weights derived from tract tracing in five connectomes: the {29 × 29}, {29 × 91}, {40 × 40}, and {40 × 91} versions of the Markov parcellation, with the number of target and source regions, respectively, indicated in each case (Markov et al, 2012; Froudist-Walsh et al, 2021), and the whole cortex connectome based on the separate regional mapping (RM) parcellation (Markov et al, 2012; Froudist-Walsh et al, 2021; Shen et al, 2019). Asterisks indicate significantly increased correlation with tract tracing data for MIND networks compared to MSNs, determined by bootstrapping the edge weights: *0.01 < *P* < 0.001, \*\**P* < 0.001. B) Correlation between tract tracing {40 × 40} weights and MIND network or MSN edge weights over a range of tract tracing network densities. C) Scatterplot of the correlations between tract-tracing weights and MRI similarities for the set of edges connecting each regional node to the rest of the connectomes; thus each point represents the correspondence between tract tracing weights and structural similarity for each region in the {40 × 40} connectome (averaged for afferent and efferent connections, see Methods for details). The dashed line *y* = *x* highlights that similarities estimated by MIND were generally more strongly correlated with tract-tracing weights (above the line of identity) than morphometric similarities. D) Radar plots of the stability of the correlation between axonal connectivity, again from the {40 × 40} connectome, and structural similarity from MSNs or MIND networks, estimated over all possible input feature sets with one or two missing features. Missing features are noted at each radial position, with the radius from the center indicating correlation with tract-tracing weights. Best case correlations for each type of structural similarity network estimated using all 6 MRI features are shown as dashed lines: SD, sulcal depth; MY, myelination (T1/T2 ratio); MC, mean curvature; SA, surface area; Vol, grey matter volume; CT, cortical thickness. Shading for A-B represents 95% CI.

Replicating and extending the work by Seidlitz et al (2018), which used a different macaque MRI dataset, we found that edge weights of axonal connectivity estimated from tract-tracing data were positively correlated with the corresponding edge weights of structural similarity estimated from MRI data by MSN or MIND network analysis (Fig. 3A). Axonal connectivity weights were significantly more positively correlated with MIND network edges than with MSN edges across all five connectomes analyzed (*P* < 0.01 from edge bootstrapping, Bonferroni-corrected). Using the {40 × 40} matrix (the largest weighted connectome with complete source and target data), we recapitulated this result over a range of tract-tracing network densities (Fig 3 B). Moreover, the degree to which regional profiles of MIND and MS corresponded to a region’s tract tracing connections was highly correlated (*r* = 0.78), though MIND showed a higher correspondence with regional tract-tracing for 85% of regions (Fig. 3C).

To test the contribution of individual morphometric features, we recalculated the correlations between the {40 × 40} tract-tracing connectome and structural similarity networks estimated with all possible subsets of four or five (of the total set of six) MRI features. The greater positive correlation of tract-tracing with MIND networks, compared to MSNs, was maintained across all feature subsets (Fig. 3D).

### Transcriptional similarity and structural similarity networks

The finding that morphometric similarity networks are spatially co-located with transcriptional similarity or gene co-expression networks (Seidlitz et al, 2018) has spurred subsequent research efforts to link MRI-derived connectomes to underlying transcriptional patterns (Seidlitz et al, 2020; Morgan et al, 2019; Zhang et al, 2021).

Following standardized processing protocols (Arnatkevičiūtė et al, 2019), we combined high-resolution spatial gene expression data on six post-mortem adult donors from the Allen Human Brain Atlas (AHBA) to generate an expression matrix for 15,633 genes in 34 regions from the left hemisphere of the DK atlas (Hawrylycz et al, 2012, 2015; Markello et al, 2021). We then calculated the pair-wise similarity of regional expression profiles to generate a {34 × 34} matrix of transcriptional similarity.

MIND networks (parcellated in the DK template) demonstrated a remarkably strong correspondence with the brain transcriptomic co-expression network (Figure 4). At the edge level, there was a greater than three-fold increase in correlations between edge weights of transcriptional similarity and MIND networks (Pearson’s *r* = 0.76, Spearman’s *ρ* = 0.81) compared to the equivalent correlations for MSNs (*r* = 0.23, *ρ* = 0.23). At the nodal level, there was an approximately two-fold increase in correlations between weighted degrees of transcriptional similarity and MIND networks (*r* = 0.85, *ρ* = 0.88), compared to the equivalent correlations for MSNs (*r* = 0.47, *ρ* = 0.30). A similar result was obtained when including the mean regional grey matter volume as a covariate (*r* = 0.75 for MIND networks, *r* = 0.5, for MSNs), suggesting results were not driven by mean volume.

**Fig. 4.**
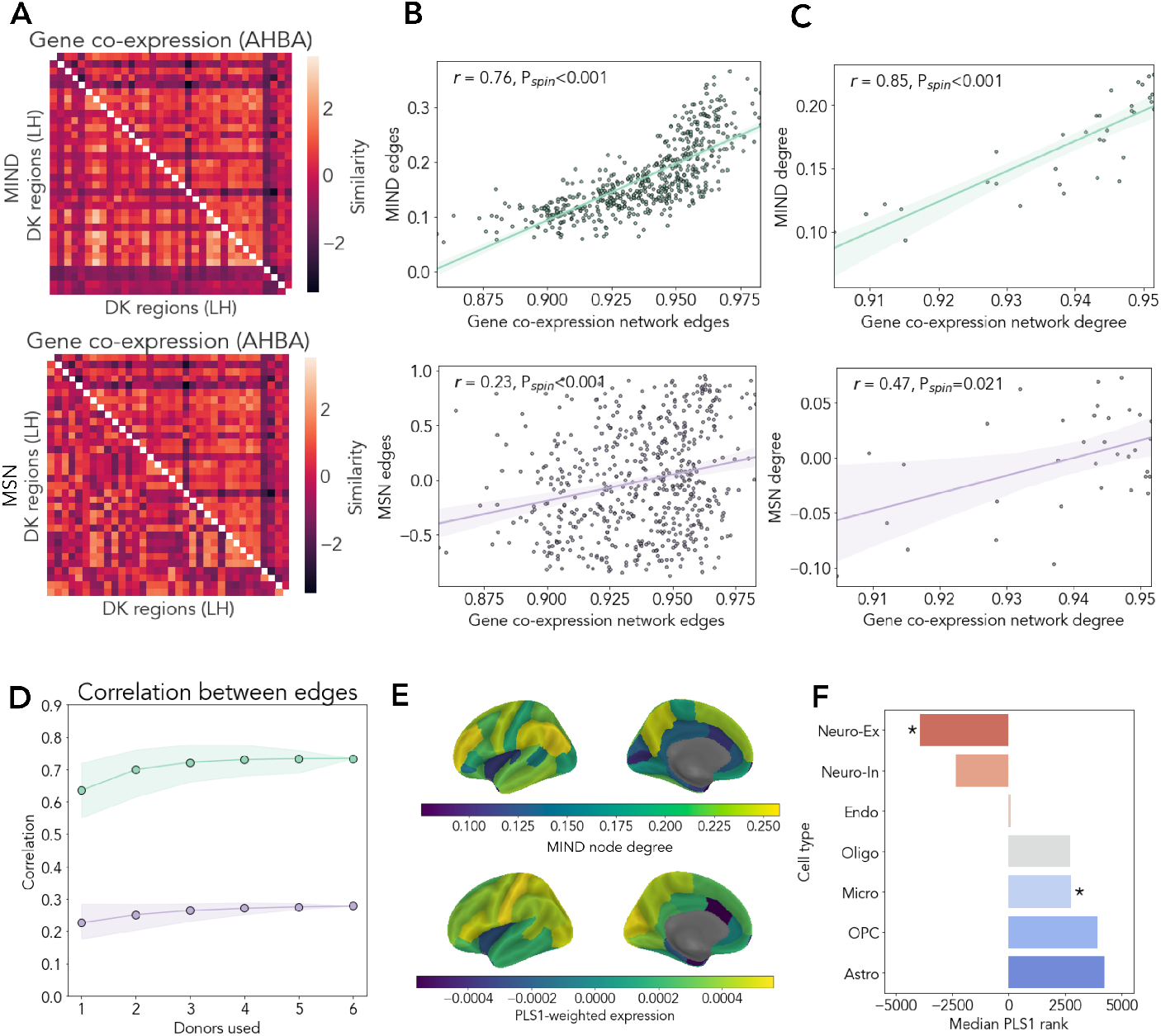
Structural similarity and transcriptional co-expression networks. A) Gene co-expression networks (upper triangles) compared to MIND networks and MSNs (lower triangles). B-C) Correlation between the gene similarity network and the group MSN or MIND network, at the level of B) edges and C) weighted regional degrees. Significance was measured using a ‘spin’ test to correct for spatial autocorrelation (see Methods). D) Stability of the correlation between structural and transcriptomic networks constructed from subsets of the 6 post-mortem brain gene expression datasets available. For each number of donors included, all combinations of transcriptional networks were constructed (without gene filtering) and the mean edge correlation was calculated. There were 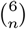 possible networks created for *n* = 1, 2 … 6 included donors. Shading indicates the minimum and maximum value of the association observed for each number of included donors. E) Cortical brain maps of MIND weighted node degree beside a weighted gene expression map derived from partial least squares analysis (PLS) of the covariation between degree, or “hubness”, of MIND nodes and gene expression. The first PLS component (PLS1) explained a significant amount of covariance (62%, *P_spin_* = 0.01) between these two modalities. F) Cell-type enrichment of the weighted, ranked gene list from PLS analysis of covariation between MIND degree and gene expression, using the median loading rank within one of seven sets of genes, each characteristic of a canonical class of cells in the central nervous system: excitatory neurons (Neuro-Ex), inhibitory neurons (Neuro-In), endothelial cells (Endo), astroctyes (Astro), microglia (Micro), oligodendrocytes (Oligo), and oligodendroglial precursor cells (OPC). The zero position on the *x*-axis represents the median position of all 15,633 genes (position 7,816), with negative ranks indicating genes that have expression positively correlated with MIND node degree, i.e., over-expressed at highly-connected MIND regions (“hubs”). Asterisk (*) indicates FDR-corrected *P* < 0.05 after a permutation test to correct for both spatial autocorrelation in the brain and correlation structure in gene expression (see Methods).

We test the robustness of the strong relationships between MIND measures of structural similarity and transcriptional similarity through several sensitivity analyses: (i) constructing different transcriptional similarity networks based on all possible subsets of 6 donor brains (Fig. 4D, S6); (ii) changing the gene inclusion criteria based on varying thresholds of differential stability (Hawrylycz et al, 2015); and (iii) replicating these analyses based on the DK template with the finer-grained DK-318 cortical parcellation (Fig S7). Under all conditions, we found that MIND network edge weights and weighted degrees remained strongly correlated with edge weights and weighted degrees of anatomically commensurate transcriptional similarity networks.

#### Cell-type specific transcriptional profiles and MIND network degrees

To characterize the relationship between MIND degree and cell-typical gene expression, we used partial least squares (PLS) to relate the {15,633 × 34} matrix of regional gene expression with the {34 × 1} vector of group-averaged MIND network weighted degree. The first PLS component (PLS1) explained a significant amount of covariance (62% variance explained, *P_spin_* = 0.01, using a “spin” permutation test to correct for cortical spatial autocorrelation as per Váša et al (2017)). Figure 4E shows the similarity between MIND degree and the cortical map of PLS-aligned transcription, calculated by averaging the spatial expression of all genes weighted by their PLS1 loadings.

Using published lists of genes specific to neuronal and glial cell types (Seidlitz et al, 2020), we calculated the median rank of genes in the PLS1 loadings within each cell-typical gene set, in line with prior enrichment work (Dorfschmidt et al, 2022; Seidlitz et al, 2020; Morgan et al, 2019). PLS1 was positively enriched for neuronal genes and negatively enriched for glial genes, with significant enrichment found for excitatory neurons and microglia (Figure. 4 F). The result that MIND network hubs were located in cortical areas with high levels of neuron-typical transcription was consistent with the observation that MIND network degree was correlated with axonal connectivity in the macaque brain, given existing work demonstrating both higher tract-tracing connectivity between transcriptionally-similar brain regions in mice (French and Pavlidis, 2011), and increased likelihood of connectivity between neurons with similar transcriptional profiles in *C. elegans* (Arnatkevičiūtė et al, 2018).

### Heritability of structural similarity network phenotypes

To characterize the extent of genetic influences on structural similarity networks, we first estimated the twin-based heritability 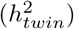 for each of the 5 MRI features measured at each region, and for each edge weight and weighted degree of the MSNs and MIND networks derived from them. Using 641 twin pairs (366 dizygotic, 275 monozygotic, total *N_twins_* = 1, 282) from the ABCD cohort, we fitted a standard ACE model to estimate additive genetic (A), shared environmental (C), and unique environmental (E) components of variance and to estimate twin heritability for each phenotype (see Methods).

MIND demonstrated increased twin-based heritability compared to MSNs in terms of both edge weights (mean 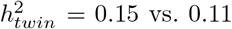, two-sided *t*-test, *P* < 0.001) and weighted nodal degree (mean 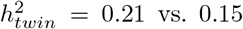, two-sided *t*-test, *P* < 0.001; Fig. 5A). To ensure the higher heritability of MIND network phenotypes compared to MSNs was not due to differing relationships with brain size (see Figure S5 for details), we confirmed that MIND network degree demonstrated increased twin-based heritability than MSN degree (two-sided *t*-test, P<0.001) after controlling for estimated total intracranial volume (eTIV).

**Fig. 5.**
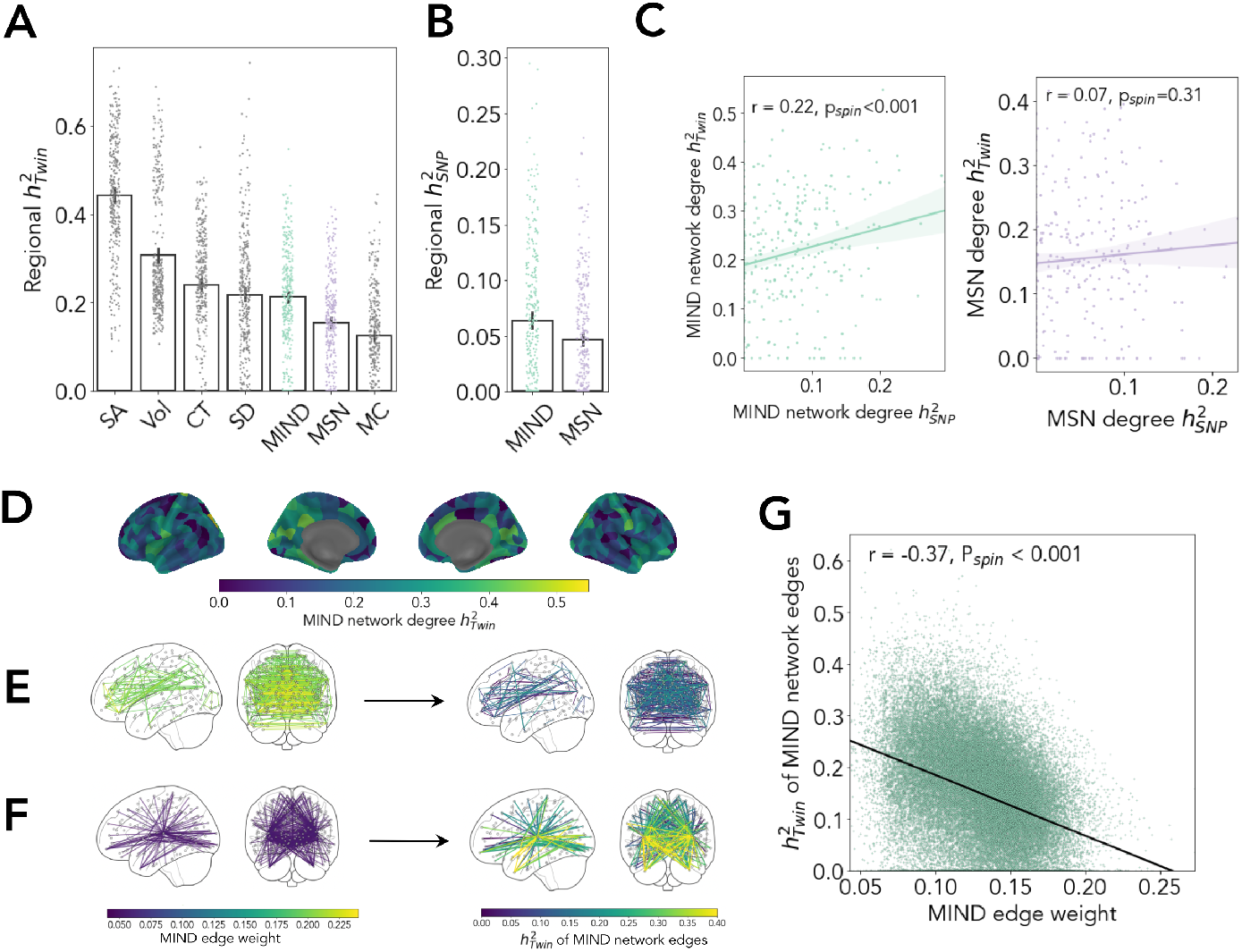
Estimating heritability, *h*^2^, of 5 regional MRI metrics and structural similarity network phenotypes derived from them. A) Twin-based heritability 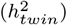 of regional MRI metrics (SA, CT, Vol, MC, and SD) and of weighted nodal degree for MIND networks and MSNs; each point represents one of 318 cortical areas. B) SNP-based heritability 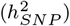 for weighted degree of MIND networks and MSNs. C) Scatterplot of twin-based *versus* SNP-based *h*^2^ estimates for weighted degree of MIND networks and MSNs; each point represents a regional node in the cortical network. D) Cortical map of the regional 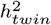 for MIND network degree. E-F) The strongest (E) and weakest (F) 1% of MIND edges and their corresponding 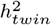 estimates. G) Scatterplot of 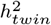 *versus* MIND network edge weights, with fitted line indicating significant negative correlation; each point is an edge in the network. For A-B), confidence intervals indicate standard error of the mean.

The five regional MRI features had average twin-based heritabilities ranging from 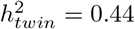 for surface area to 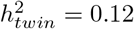 for mean curvature. The average heritability MIND weighted degree was significantly higher than 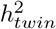 for regional mean curvature (two-sided *t*-test, P<0.001), comparable to the 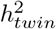 for regional estimates of mean sulcal depth (two-sided *t*-test, P>0.05), and lower than the heritabilities of the three macro-structural MRI metrics related to the size of each regional node of cortex (surface area, cortical thickness, and volume; two-sided t-tests, all *P* < 0.001). The cortical maps of regional MRI heritability for the different MRI features were positively correlated with each other (0.09 < *r* < 0.61; Fig S8). This result points to the existence of a general anatomical gradient of the heritability of brain structure, where similar patterns of heritability are observed across different MRI phenotypes.

#### SNP-based heritability

We estimated SNP-based heritability for weighted degree in MSN and MIND networks using genetic data from 4,085 *unrelated* individuals of predominantly European genetic ancestries from the ABCD cohort; and GCTA (Yang et al, 2011) software for genome-wide complex trait analysis.

SNP-based heritability for weighted degree of MIND networks (mean 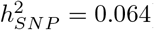) was greater than for degree of MSNs (mean 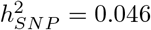), and this difference was significant (two-sided *t*-test *P* < 0.001; Fig. 5B). SNP-based and twin-based heritabilities were positively correlated for weighted degree of MIND networks (*r* = 0.22, *P_spin_* < 0.001), but were not correlated for degree of MSNs (*r* = 0.07, *P_spin_* = 0.31; Fig. 5C). This demonstrates that common genetic variants partly explain variance in MIND networks.

#### Increased heritability of MIND between structurally divergent regions

Twin-based heritabilities for MIND network edges were robustly, negatively correlated with edge weights (*r* = −0.37, Fig. 5G). This is visualized in Fig. 5 E-F, where the highest MIND edges, between most similar areas of cortex, e.g. inter-hemispheric connections, have much lower heritability than the lowest MIND edges, between most dissimilar areas of cortex, e.g., connections between neocortical areas and areas of insular and limbic cortex. We observed no correlation between Euclidean distance and edge heritability (*r* = 0.02, *P_spin_* = 0.66), despite an exponentially decaying relationship between distance and MIND edge strength (Fig. S3).

MIND network weighted degree was also negatively correlated with heritability (*r* = −0.24, *P_spin_* = 0.02). When categorized by cytoarchitectonic class (Fig. S8), weighted degree was more strongly heritable (mean 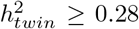) for insular, primary sensory, and limbic cortex; and less strongly heritable for primary motor, association and secondary sensory cortex (mean 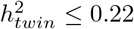). The difference in heritabilities between cytoarchitectonic classes was significant (ANOVA, *F*_6,311_= 7.54; *P* < 0.001). Thus MIND network “hub” nodes that have a high degree of similarity with many other nodes in the network, and are typically located in motor and association cortices, were less strongly heritable than “non-hub” nodes that have a high degree of dissimilarity with many other cortical areas, and are typically located in insular, primary sensory, and limbic cortex (Fig. S8).

## Discussion

Here we introduced morphometric inverse divergence (MIND) as a novel metric of structural similarity between cortical areas and showed that MIND networks have greater reliability and biological validity compared to morphometric similarity networks (MSNs) derived from the same set of MRI features in a large sample. At a technical level, the relative superiority of individual brain connectome mapping by MIND networks compared to MSNs is simply explained. MIND measures similarity by the divergence between multi-dimensional distributions with many degrees of freedom, whereas MSNs are predicated on regional summary statistics of each MRI feature and are therefore less efficiently estimated on many fewer degrees of freedom.

This fundamental difference between MIND and MSN estimators of structural similarity greatly enhanced the reliability and validity of the resulting MIND networks. First, MIND network edge weights and nodal degrees were more consistent between subjects, more resilient to the inclusion of noise features, and more robust to different parcellation templates used to define cortical nodes. The greater parcellation consistency of MIND networks makes intuitive sense: if the boundaries of a particular region change slightly between atlases, the multi-dimensional distribution of MRI features at the vertex-level will remain relatively consistent, whereas MSNs are more sensitive to parcellation differences because even small shifts in areal boundaries will have marked effects on macro-structural metrics, such as volume or surface area, which were included in the short feature vector used to estimate the morphometric similarity between regions.

Benchmarking both MIND networks and MSNs against prior principles of cortical network organization (von Economo and Koskinas, 1925; Barbas, 2015; Goulas et al, 2016), we found that MIND networks were more representative of symmetrical inter-hemispheric connections, connections between areas belonging to the same cytoarchitectonic class, and between areas with axonal inter-connectivity demonstrated by the gold standard of retrograde tract-tracing in the macaque monkey. These results consistently indicate that the connectomes rendered by MIND analysis of structural similarity are more aligned with the principles that structural similarity between regions should be greater for bilaterally homologous cortical areas, for cytoarchitectonically homogenous areas, and for axonally connected areas.

Recent work has begun to establish another principle of brain network organization, that structurally similar or axonally inter-connected regions will typically have more similar profiles of gene transcription than cytoarchitectonically dissimilar or unconnected pairs of regions (Arnatkevičiūtė et al, 2021). In short, the structural architecture of the connectome recapitulates the organisation of the brain gene co-expression network. We therefore expected, and confirmed, that the more reliable and valid connectomes produced by MIND analysis should be more strongly correlated than MSNs with a gene co-expression network derived from the Allen Human Brain Atlas. The significantly greater strength of association between transcriptional similarity and structural similarity measured by MIND was clearly evident at the level of both edges and nodes. Moreover, the high degree hubs of MIND networks were significantly co-located with areas where neuron-specific genes were highly expressed (Seidlitz et al, 2020). These results strongly support the preferred use of MIND network analysis for future imaging studies designed to discover the transcriptional mechanisms underpinning anatomical connectomes in health and disease.

However, the causal relationship(s) between MIND metrics of structural similarity and gene co-expression metrics of transcriptional similarity are not resolved by these correlational results. Several causal pathways could explain the strong coupling between MIND and transcriptional networks. Spatially patterned and developmentally phased gene expression drives the expansion and development of the human cortex (Geschwind and Rakic, 2013), so it is at least plausible that the network organization of transcription is an important driver or template of the network organization of the structural similarity and axonal connectivity of the cortex.

To probe the biological drivers of MIND more directly, we took the first steps towards a genetic analysis of MRI similarity network phenotypes. We demonstrated that MIND network phenotypes have higher twin-based and SNP-based heritabilities than comparable MSN phenotypes, further endorsing the biological validity of MIND networks compared to MSNs and setting the scene for future, more detailed investigation of genetic effects on MIND network phenotypes.

It is notable that the heritability of the MIND similarity between two regions was found to be higher for edges between structurally dissimilar or differentiated regions, e.g., connecting limbic, insular or primary sensory cortical areas to the rest of the network. Consequently, MIND network hubs in motor and association cortex, with a high degree of similarity to many other neocortical areas, were less genetically influenced than primary sensory, insular and limbic non-hub regions with more distinctive, less generally similar cytoarchitecture. These results may reflect the notion that the heritability of brain phenotypes is inversely related to plasticity (Vainik et al, 2020; Haak and Beckmann, 2019), with *decreased* heritability reflecting *increased* plasticity observed for motor and association areas rather than more conserved primary sensory and allocortical regions. This interpretation is also consistent with the finding of relaxed genetic control over cortical organization of recently-evolved association cortex regions relative to other primates (Gómez-Robles et al, 2015).

One limitation of MIND networks – shared by MSNs – is that they do not currently include subcortical regions, which are not represented by surface reconstructions. Moreover, while MIND networks are heritable and show a strong relationship to transcriptional networks, it remains unclear whether the genetic influence in MIND is mediated by changes in cortical gene expression. Future work in the form of a large-scale GWAS, perhaps complemented with analysis of data containing both *in vivo* imaging and postmortem cortical expression data (Bennett et al, 2018), will be necessary to identify the genetic loci and genes associated with changes in MIND network phenotypes. Ultimately, we expect the study of MIND networks to provide a new perspective on the principles of cortical organization that reflect the genetic architecture of the brain and which may underlie human brain development, aging, and disease.

## Methods

### Data and code availability

Python code for MIND calculation using standard FreeSurfer outputs is publicly available at: https://github.com/isebenius/MIND.

The preprocessed macaque data (Xu et al, 2020; Milham et al, 2018) can be accessed at https://balsa.wustl.edu/reference/976nz. Tract-tracing connectomes based on the Markov parcellation can be accessed through https://core-nets.org. The multimodal connectome using the RM parcellation (as well as the RM atlas itself) can be accessed at https://zenodo.org/record/1471588#.YqBt5S2ca_U. Data from the ABCD cohort requires access to the NIMH data archive (NDA) and can be applied for at https://nda.nih.gov/abcd.

### MIND estimation

#### Definition of the MIND similarity metric

Here we describe the definition of MIND as a statistical metric of structural similarity given a surface reconstruction of the cortex. This surface can be described by a set of vertices 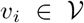, where each *v_i_* is a vector of d structural features such as cortical thickness and sulcal depth. These features (interchangeably described as structural and morphometric features) are automatically generated at the vertex-level by FreeSurfer’s `recon-all` command. A cortical parcellation with *R* regions is a partition of 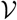 such that 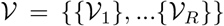. For each region *r*, we let *P_r_* be the true multivariate distribution of structural features from which 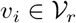 are observations.

For a given pair of regions *a* and *b*, we estimate *D*_*KL*_(*P_a_* || *P_b_*), the Kullback Leibler (KL) divergence between them. Because KL divergence is not symmetric, we use a commonly-used symmetric version of the metric, computed as follows and in line with previous work (Homan et al, 2019; Wang et al, 2016):

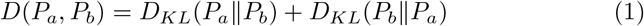

We define the Morphometric Inverse Divergence (MIND) similarity metric, bounded between 0 and 1, as follows:

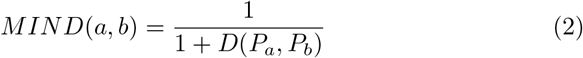

#### Multivariate Kullback Leibler divergence estimation

One key challenge in calculating MIND networks is appropriately estimating multivariate KL divergence. Traditionally, KL divergence between empirical distributions is calculated by a two-step approach: i) non-parametric estimation of the probability density functions (PDFs) of the observed data; and ii) computing the divergence using the approximated PDFs. However, the initial density estimation step of this approach is sensitive to many choices of parameters (Leming et al, 2021; Homan et al, 2019; Wang et al, 2016; Perez-Cruz, 2008). Extended to multiple dimensions, density estimation becomes especially problematic; for multivariate data with as few as three dimensions, standard non-parametric density estimators provide very poor results (Wang and Scott, 2019). While research into alternate methods for higher-dimensional density estimation is actively ongoing, no consensus currently exists on an effective, efficient method for this purpose (Wang and Scott, 2019).

Here, we circumvent the need to perform the difficult first step of density estimation by leveraging a *k*-nearest-neighbor approach (Perez-Cruz, 2008) for calculating multivariate KL divergence directly from the observed vertex-level data. This approach has significant advantages compared to explicit density estimators: namely, it does not require the specification of any parameters and it can be computed efficiently (Brown, 2014).

More formally, given regions *a* and *b*, vertices *V_a_* and *V_b_*, with true multivariate distributions *P_a_* and *P_b_*, the KL divergence between *P_a_* and *P_b_* is defined mathematically as follows:

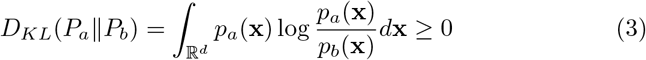

We used the *k*-NN divergence approximation by Perez-Cruz (2008) to estimate *D*_*KL*_(*P_a_*||*P_b_*):

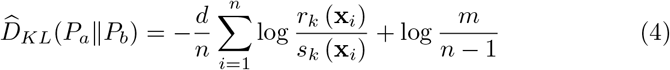

Here, *d* is the number of structural features used, 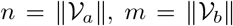, and *r_k_*(**x**_*i*_) and *s_k_*(**x**_*i*_) are the Euclidean distances of **x**_*i*_ to the *k*-th most similar vertex in of **x**_*i*_ in 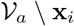 and 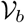, respectively. We use *k* = 1 (nearest-neighbor) in our analysis, and we calculate *r_k_* (**x**_*i*_) and *s_k_* (**x**_*i*_) efficiently using K-D trees, a method for data representation that enables the rapid lookup of nearest neighbors.

To account for the unlikely but possible occurrence that the estimation of KL is negative, we set the minimum value of 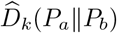 to be zero. A symmetric measure of KL divergence was then given by:

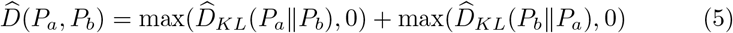

And MIND was finally estimated by:

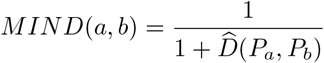

#### Standardizing and filtering vertex-level data

Because each feature is measured on different scales, we standardize (Z-score) each feature across all vertices in the brain before before parcellating the data into vertex-level distributions and calculating MIND.

Additionally, structural vertex-level data can sometimes represent biologically-unfeasible conditions; namely, when vertices have values of zero for cortical thickness, volume, or surface area. For MIND estimation, we discarded all such vertices. One result of this filtering step is that if a region is left with zero or one vertices, a complete MIND network cannot be computed. Thus, using parcellations of smaller parcel size will generally lead to higher likelihood that one or more regions contains no vertices, and therefore fewer networks that can be fully calculated. Removing the condition that all vertices must have above-zero values of thickness, volume, and area will mitigate this, though at the trade-off of including vertices that correspond to potentially unfeasible conditions.

#### Computational costs of MIND network analysis

Construction of MIND networks can be completed at reasonable computational cost. Given *d* dimensions and *n* vertices, a K-D Tree can be constructed in 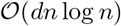 – with more recent approaches further improving a worst-case construction time to 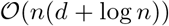 (Brown, 2014) – and can be queried in 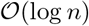 (Buitinck et al, 2013; Bentley, 1975). This computation is not a bottleneck; in practice, we observed that the computational resources expended during structural image (FreeSurfer) preprocessing far exceeded MIND network computation. For reference, on consumer hardware, computation of a single MIND network ranged from roughly 1 minute (68-ROI DK atlas) to roughly 10 minutes (360-ROI HCP atlas).

#### MSN calculation

MSNs were computed as described in Seidlitz et al (2018). Specifically, we considered the widely-used summary statistics computed by FreeSurfer’s `mris_anatomical_stats` command to characterize each region. The five summary statistics describing each region are the following:

- Mean sulcal depth
- Mean cortical thickness
- Total volume
- Total surface area
- Integrated rectified mean curvature

Each features was Z-scored, and the resulting MSN was defined as the pairwise Pearson correlation between all vectors of the five standardized features. To construct MSNs with 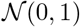 noise features (as in Fig. 2F), the Gaussian noise columns were added as new features at each vertex, averaged within each region, and Z-scored across regions before inclusion into the vector of structural features.

### Human MRI datasets

The Adolescent Brain Cognitive Development (ABCD) cohort currently comprises T1-weighted structural MRI data on 11,449 participants (including 697 twin pairs) aged 9-11 years at baseline scanning. Recruitment for the ABCD study was intended to generate a diverse, representative sample (Garavan et al, 2018) for the longitudinal study of brain development and cognition. This work is registered as study #1796 on the NIMH data archive, DOI 10.15154/1528079. The number of subjects included for different analyses can be visualized in Fig. S2. After data filtering and quality control (QC), this led to to 10,367 subjects included in our principal dataset parcellated in DK-318 (N=10,353 for DK, and N=9,218 for HCP parcellation, with the difference between parcellation due to the vertex filtering step described above). Group-level MIND networks and MSNs were constructed from these cohorts. These 10,367 subjects in DK-318 parcellation included 641 complete twin pairs (366 dizygotic, 275 monozygotic), which served as the cohort for estimating twin-based heritability. To estimate SNP-based heritability, we used a sample of 4,085 subjects, comprising unrelated participants of European ancestry with MRI and genetic data that passed QC criteria. To study the effect of including varying numbers of noise columns (Fig. 2F), we used a random subset of 150 subjects to avoid the cost of constructing many network versions for all 10,367 individuals.

Extensive documentation of the scanner types and protocols used for MRI in the ABCD study can be found in Hagler et al (2019); Casey et al (2018). T1-weighted images were 1 mm isotropic, RF-spoiled gradient echo using prospective motion correction if available, and from one of three (3T) scanner models: Siemens (Prisma VE11B-C), Philips (Achieva dStream, Ingenia), or GE (MR750, DV25-26) (Fischl et al, 1999; Hagler et al, 2019). The images were processed using FreeSurfer Version 5.3.0.

#### MRI quality control

To ensure high quality of the included scans, we used the Euler number (Fischl, 2012), an index of scan quality generated automatically by FreeSurfer. Figure S2 shows the distribution of Euler number in the entire ABCD cohort, with some extreme outliers in the sample. To discard these scans, we used a cutoff threshold of −120 corresponding to a median absolute deviation (MAD) score of ≥2.6.

#### Site-related batch effects

ABCD is a multi-site study with well-known batch effects due to scanning at different sites (Nielson et al, 2018). To correct these site-related batch effects in quality-controlled data, prior to MIND analysis, we used NeuroCombat v0.2.12 (Fortin et al, 2018), an adaptation of the standard ComBat batch correction tool (Johnson and Li, 2007) designed specifically for structural MRI brain data. While adjusting for site-specific effects in this manner, we included age (in months) and sex to be biologically-relevant covariates (i.e. differences in age and sex distribution between sites were not considered site-specific effects).

#### Cortical parcellation

We used the Desikan-Killiany (Desikan et al, 2006), DK-318 (Romero-Garcia et al, 2012), and HCP (Glasser et al, 2016) parcellations in this work. To compare edge-wise consistency between group-level DK and DK-318 atlases (Figure 2H), we assumed that an edge between two regions in the DK atlas should be comparable to that derived by averaging all edges between the sub-divisions of the regions in the finer-grained DK-318 atlas. In this manner, we interpolated the group DK-318 networks back to the original DK atlas, then compared these recreated networks to those computed directly on the original 68-region DK parcellation. The correlation between the original DK and interpolated (DK-318 interp.) networks was used to measure edge-level parcellation consistency.

The mapping from each region in the 318-region subdivision of the Desikan Killiany atlas (Fig. 2I) was based on the mapping used in Seidlitz et al (2018), originally performed by Vértes et al (2016) and Whitaker et al (2017). These prior studies used the closely related DK-308 parcellation (Romero-Garcia et al, 2012), which is an asymmetric version of the DK-318 atlas. We used a simple majority-voting procedure to translate the DK-308 parcellation to the related DK-318 atlas.

### Twin-based heritability

We used the `umx` package version 2.10.0 to implement a structural equation model (SEM) of the ACE model to infer heritability estimates (Bates et al, 2019; Neale and Cardon, 1994). The ACE model estimates the contributions to observed variance due to additive genetic (A) in the context of common (C) and specific environmental (E) effects on variance (Bates et al, 2019; Neale and Cardon, 1994). We defined heritability (*h*^2^) as the proportion of variance due to additive genetics contributions such that 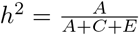 (Verhulst et al, 2018).

The SEM model does not estimate A, C, and E directly, but rather estimates path coefficients *a*, *c*, and, *e* such that *a*^2^ = A, *c*^2^ =C, and *e*^2^ = E. These path coefficients are sometimes directly used as reports of heritability (e.g. Bethlehem et al (2022)). To validate our processes for twin-based heritability estimation, we replicated a previously reported estimate of 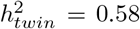 for whole-brain grey matter volume (GMV) in the ABCD cohort (Bethlehem et al, 2022). A discussion comparing our work to previous heritability estimates using path coefficients can be found in the “Additional heritability analyses” section of the Supplementary Materials.

### Human genetic data QC and SNP-based heritability

Full details of genetic quality control procedures are provided by Warrier et al (2021, 2022a). Briefly, we excluded SNPs with genotyping rate < 90%, and individuals with genotyping rate < 95%, whose genetic sex did not match their reported sex. We identified individuals of predominantly European genetic ancestries using multidimensional scaling after including samples from the 1000 Genomes phase 3 data (Fairley et al, 2019). In the subset of individuals of predominantly European ancestries, we further exlcuded SNPs not in Hardy-Weinberg equilibrium (*p* < 1E − 6) and individuals with excessive heterozygosity. Related individuals (> 5% identity by state) were excluded using the Genome-wide Complex Trait Analysis Genome-based REstricted Maximum Likelihood (GCTA-GREML) software (Yang et al, 2011) prior to estimation of SNP-based heritabilities using a genetic relatedness matrix. The genetic relatedness matrix was derived from genotyped samples after controlling for age, age^2^, age×sex, age^2^×sex, sex, imaging centre, mean framewise displacement, maximum framewise displacement, Euler Index, and the first ten genetic principal components as covariates.

### Macaque MRI and tract-tracing data

We used MRI data from 19 female rhesus macaque monkeys (*Macaca mulatta*; aged 18.5-22.5 years) in the UC-Davis cohort provided by the PRIME-DE resource (Milham et al, 2018). The animals were anesthetized and scanned on a Siemens Skyra 3T MRI with a 4-channel clamshell coil with 0.3 isotropic resolution (TR = 2500ms) (Milham et al, 2018). These data were pre-processed using the HCP Non Human Primate (Autio et al, 2020) pipeline by Xu et al (2020).

All individual scans were previously spatially coregistered with the group-level Yerkes19 atlas (Donahue et al, 2016). On this basis, we constructed group-level structural similarity networks by first averaging vertex-level features and then constructing a MIND network and an MSN. We calculated MSNs by manually generating the same output as performed by FreeSurfer’s `mris_anatomical_stats` command used to calculate human MSNs. Specifically, within each region, vertex values of sulcal depth and thickness were averaged, volume and surface area were summed, and mean curvature was absolute-valued, multiplied by surface area, and summed (thus outputting integrated rectified mean curvature).

We used four tract-tracing connectivity matrices based on the 91-region Markov M132 parcellation of the left hemisphere (Markov et al, 2012): the original and most widely used {29 × 29} complete connectivity matrix between 29 cortical areas used both as source and target regions in retrograde tract-tracing experiments; the {29 × 91} matrix including all originally measured source-target connections (Markov et al, 2012); and the corresponding {40 × 40} and {40 × 91} matrices from a recently published extension of the original Markov dataset which increased the number of target regions from 29 to 40 (Froudist-Walsh et al, 2021). For all these tract-tracing connectomes, we used the log-transformed fraction of labeled neurons (log(FLNe)) as the measure of axonal connectivity (Markov et al, 2012; Seidlitz et al, 2018). Additionally, we used a bihemisperhic connectivity matrix based on the independent Regional Mapping (RM) parcellation and estimated by using diffusion weighted imaging to infer the connectivity weights from categorical estimates derived from the CoCoMac database (Shen et al, 2019; Bakker et al, 2012).

To study the relationship between structural similarity and regional connectivity profiles, we generated two tract-tracing connectivity profiles per region based on the vectors of afferent and efferent edges connected to a node. We correlated both of these vectors with the node’s (undirected) profile of structural similarity (MIND or MS), Fisher transformed the two correlations, averaged, and inverse transformed in order to calculate a final correlation between a region’s tract-tracing connectivity and structural similarity (reported in Figure 3C).

To test for the difference between the correlation between MIND networks and tract-tracing versus that between MSNs and tract-tracing, we performed edge-wise bootstrapping with the one-sided null hypothesis that MIND did not have a greater correlation than MSNs with tract-tracing. Thus for each bootstrapped edge sample, we calculated the difference between the tract-tracing correlation for MIND and MSNs, then calculated the P value as the fraction of samples for which this value fell below zero. We used a significance threshold of *α* = 0.01 corresponding to a Bonferroni correction for the five connectomes used.

### Gene expression analysis

#### Data and pre-processing

The Allen Human Brain Atlas (AHBA) contains high-resolution spatial genome transcriptional data in the cortex from 6 post-mortem brains (male/female = 4/2, mean age = 45 years). We focused our analysis of the Allen Human Brain Atlas on the Desikan-Killiany (DK) parcellation of human brain gene expression maps, given prior work on standardizing the pre-processing pipeline for this atlas (Arnatkevičiūtė et al (2019); Markello et al (2021); Hawrylycz et al (2015)) and because the coarse-grained DK parcellation ensures high donor coverage for all regions. Only 2 brains provided data from the right hemisphere, so we focused on the left hemisphere only and we used the abagen package (v0.1.3) developed by Markello et al (2021) with default settings to fetch (the `get_expression_data` command) and manipulate the AHBA data. These pre-processing steps included aggregating probes across all available donors, selecting probes using an intensity-based filtering threshold of 0.5, normalizing microarray expression values for each sample and donor using the scaled robust sigmoid function, and combining gene probes for each region within each donor before combining across donors.

#### Transcriptional similarity metric

We used an angular similarity metric based on cosine distance (rather than raw cosine similarity or pearson correlation) to measure transcriptional similarity between regions. If *g_x_* and *g_y_* are the vectors of gene expression for regions X and Y, the transcriptional similarity between the two regions is defined as:

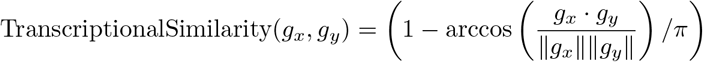

The choice of this metric was informed by the prior finding (Cer et al, 2018) that it more precisely measures differences between high-dimensional vectors with high average similarity, which is the case for regional transcription data. In this formulation, ||·|| the operator represents the length of the vector (the square root of the sum of squares of all elements). In the case of no gene filtration, all 15,633 genes are used for each region.

#### Cell-type specific gene enrichment analysis

To perform enrichment analysis of the genes most highly co-located with MIND degree, we first performed a partial least squares (PLS) regression between the {15,633 × 34} matrix of AHBA gene expression and the {34 × 1} vector of MIND weighted node degree, then ranked each gene based on their position in the list of PLS1 loadings, with lower rank corresponding to genes with higher positive correlation with MIND degree. We then leveraged the extensive meta-analysis performed by Seidlitz et al (2020) to assign 4,110 genes to one of seven major classes of cells in the central nervous system: excitatory neurons, inhibitory neurons, endothelial cells, astroctyes, microglia, oligodendrocytes, and oligodendroglial precursor cells. To measure the cell-type enrichment in the loadings of the first component of the PLS, we calculated the median rank of each set of cell-typical genes. Then, we utilized a permutation test to account for both the intrinsic correlation structure in the AHBA expression data as well as spatial autocorrelation in the cortical map of MIND degree, detailed below.

### Spin permutation tests

We adopted the widely used ‘spin’ test to measure for significance of association between two cortical maps while correcting for spatial autocorrelation (Váša et al, 2017; Alexander-Bloch et al, 2018). This test uses the (x,y,z) coordinates of each parcel to generate permutations of parcellated data that maintains its spatial embedding. We used the implementation by the `gen_spinsamples` command from the netneurotools Python package with the parameter `method` set to `hungarian`, which ensures that each index is used only once per permutation and uses the Hungarian algorithm to minimize the global cost of reassignment. The same spatial permutations were applied to the left and right hemispheres in order to maintain bilateral symmetry. When testing for significance between network edges (i.e. the relationship between MIND edge heritability and mean edge strength), we used the same permutation scheme, and simply applied spatial permutations to both node sources and targets (i.e. the rows and columns of a connectivity matrix), thus in effect rotating the entire network. All such statistical tests used were two-sided, such that the null hypothesis was that the variance explained (*r*^2^) between two cortical maps or edges was not greater than the *r*^2^ expected by chance, accounting for spatial autocorrelation. We used 1000 permutations for all tests.

In order to test for the significance of cell-typical gene set enrichment in the loadings from the PLS component between gene expression and MIND degree, we generated 1000 spin test permutations of the vector of MIND degrees, while keeping the gene expression data intact. For each spatial permutation of MIND degree, we fit a new PLS model and ensured that lower loading rank corresponded with a positive correlation with the permuted brain map. For each new model, we calculated the median gene rank within each set of cell-type specific genes. The two-sided null hypothesis of our permutation test was that the median gene rank of a cell-typical gene-set of was not significantly different from the median position of all genes (rank 7,816). We thus calculated the P-value as the fraction of all permutations for which each set of cell-typical genes had a median rank farther away from rank 7,816 than the true median rank. We FDR-corrected the resulting seven P-values. In this scheme, gene sets with median PLS rank significantly *lower* than the median position were *positively* associated with MIND degree.

## Acknowledgments

I.S. was generously supported by a Gates-Cambridge Scholarship, and by the Accelerate Programme for Scientific Discovery, funded by Schmidt Futures. J.S. was supported by NIMH T32MH019112. A.A.B and J.S. were supported by NIMH K08MH120564. V.W. was supported by St. Catharine’s College Cambridge. R.A.I.B. was supported by the Autism Research Trust. R.R.G. is funded by the EMERGIA Junta de Andalucía program (EMERGIA20 00139) and the Plan Propio of the University of Seville. T.T.M. was supported by funds from NIH T32HG010464. E.T.B. was supported by an NIHR Senior Investigator award. S.E.M. was supported by the Accelerate Programme for Scientific Discovery, funded by Schmidt Futures, and a Fellowship from The Alan Turing Institute, London (EPSRC grant EP/N510129/1). We thank Dr. Lisa Ronan for help in processing the ABCD imaging data.

Data were curated and analysed using a computational facility funded by an MRC research infrastructure award (MR/M009041/1) to the School of Clinical Medicine, University of Cambridge and supported by the mental health theme of the NIHR Cambridge Biomedical Research Centre. The views expressed are those of the authors and not necessarily those of the NIH, NHS, the NIHR or the Department of Health and Social Care.

Data used in the preparation of this article were obtained from the Adolescent Brain Cognitive DevelopmentSM (ABCD) Study (https://abcdstudy.org), held in the NIMH Data Archive (NDA). This is a multisite, longitudinal study designed to recruit more than 10,000 children age 9-10 and follow them over 10 years into early adulthood. The ABCD Study® is supported by the National Institutes of Health and additional federal partners under award numbers U01DA041048, U01DA050989, U01DA051016, U01DA041022, U01DA051018, U01DA051037, U01DA050987, U01DA041174, U01DA041106, U01DA041117, U01DA041028, U01DA041134, U01DA050988, U01DA051039, U01DA041156, U01DA041025, U01DA041120, U01DA051038, U01DA041148, U01DA041093, U01DA041089, U24DA041123, U24DA041147. A full list of supporters is available at https://abcdstudy.org/federal-partners.html. A listing of participating sites and a complete listing of the study investigators can be found at https://abcdstudy.org/consortium_members/. ABCD consortium investigators designed and implemented the study and/or provided data but did not necessarily participate in the analysis or writing of this report. This manuscript reflects the views of the authors and may not reflect the opinions or views of the NIH or ABCD consortium investigators. The ABCD data repository grows and changes over time. The ABCD data used in this report came from NIMH Data Archive Digital Object Identifier (10.15154/1528079)]. DOIs can be found at [https://dx.doi.org/10.15154/1528079.

We also thank the Allen Human Brain Atlas (AHBA) for their valuable contributions to open science. For the purpose of open access, the authors have applied a Creative Commons Attribution (CC BY) licence to any Author Accepted Manuscript version arising from this submission.

## Declarations

ETB works in an advisory role for Sosei Heptares, Boehringer Ingelheim, GlaxoSmithKline, Monument Therapeutics. A.A.B. receives consulting income from Octave Bioscience.

## Footnotes

1 Because we standardize each morphometric feature, the non-random, measured variables also have mean=0 and variance=1.

## References

Alexander-Bloch AF, Shou H, Liu S, et al (2018) On testing for spatial correspondence between maps of human brain structure and function. NeuroImage 178:540–551. https://doi.org/10.1016/j.neuroimage.2018.05.070

Arnatkevičiūtė A, Fulcher BD, Pocock R, et al (2018) Hub connectivity, neuronal diversity, and gene expression in the caenorhabditis elegans connectome. PLOS Computational Biology 14(2). https://doi.org/10.1371/journal.pcbi.1005989

Arnatkevičiūtė A, Fulcher BD, Fornito A (2019) A practical guide to linking brain-wide gene expression and neuroimaging data. NeuroImage 189:353–367. https://doi.org/10.1016/j.neuroimage.2019.01.011

Arnatkevičiūtė A, Fulcher BD, Bellgrove MA, et al (2021) Imaging transcriptomics of brain disorders. Biological Psychiatry Global Open Science https://doi.org/https://doi.org/10.1016/j.bpsgos.2021.10.002, URL https://www.sciencedirect.com/science/article/pii/S2667174321001191

Autio JA, Glasser MF, Ose T, et al (2020) Towards hcp-style macaque connectomes: 24-channel 3t multi-array coil, mri sequences and preprocessing. NeuroImage 215:116,800. https://doi.org/10.1016/j.neuroimage.2020.116800

Bakker R, Wachtler T, Diesmann M (2012) Cocomac 2.0 and the future of tract-tracing databases. Frontiers in Neuroinformatics 6. https://doi.org/10.3389/fninf.2012.00030, URL https://www.frontiersin.org/article/10.3389/fninf.2012.00030

Barbas H (2015) General cortical and special prefrontal connections: Principles from structure to function. Annual Review of Neuroscience 38(1):269–289. https://doi.org/10.1146/annurev-neuro-071714-033936

Bates TC, Maes H, Neale MC (2019) Umx: Twin and path-based structural equation modeling in r. Twin Research and Human Genetics 22(1):27–41. https://doi.org/10.1017/thg.2019.2

Bennett DA, Buchman AS, Boyle PA, et al (2018) Religious orders study and rush memory and aging project. Journal of Alzheimer’s Disease 64(s1). https://doi.org/10.3233/jad-179939

Bentley JL (1975) Multidimensional binary search trees used for associative searching. Communications of the ACM 18(9):509–517. https://doi.org/10.1145/361002.361007

Bethlehem R, Seidlitz J, White S, et al (2022) Brain charts for the human lifespan. Nature https://doi.org/https://doi.org/10.1038/s41586-022-04554-y, URL https://www.nature.com/articles/s41586-022-04554-y#citeas, https://arxiv.org/abs/https://www.nature.com/articles/s41586-022-04554-y#citeas

Brown RA (2014) Building a balanced k-d tree in (kn log n) time. CoRR abs/1410.5420. URL http://arxiv.org/abs/1410.5420, https://arxiv.org/abs/1410.5420

Buitinck L, Louppe G, Blondel M, et al (2013) API design for machine learning software: experiences from the scikit-learn project. In: ECML PKDD Workshop: Languages for Data Mining and Machine Learning, pp 108–122

Bullmore E, Sporns O (2009) Complex brain networks: Graph theoretical analysis of structural and functional systems. Nature Reviews Neuroscience 10(3):186–198. https://doi.org/10.1038/nrn2575

Casey B, Cannonier T, Conley MI, et al (2018) The adolescent brain cognitive development (abcd) study: Imaging acquisition across 21 sites. Developmental Cognitive Neuroscience 32:43–54. https://doi.org/10.1016/j.dcn.2018.03.001

Cer D, Yang Y, Kong S, et al (2018) Universal sentence encoder. CoRR abs/1803.11175. URL http://arxiv.org/abs/1803.11175, https://arxiv.org/abs/1803.11175

Desikan RS, Ségonne F, Fischl B, et al (2006) An automated labeling system for subdividing the human cerebral cortex on mri scans into gyral based regions of interest. NeuroImage 31(3):968–980. https://doi.org/10.1016/j.neuroimage.2006.01.021

Donahue CJ, Sotiropoulos SN, Jbabdi S, et al (2016) Using diffusion tractography to predict cortical connection strength and distance: A quantitative comparison with tracers in the monkey. The Journal of Neuroscience 36(25):6758–6770. https://doi.org/10.1523/jneurosci.0493-16.2016

Dorfschmidt L, Bethlehem RA, Seidlitz J, et al (2022) Sexually divergent development of depression-related brain networks during healthy human adolescence. Science Advances 8(21). https://doi.org/10.1126/sciadv.abm7825

von Economo C, Koskinas G (1925) Die cytoarchitektonik der hirnrinde des erwachsenen menschen. J. Springer

Fairley S, Lowy-Gallego E, Perry E, et al (2019) The International Genome Sample Resource (IGSR) collection of open human genomic variation resources. Nucleic Acids Research 48(D1):D941–D947. https://doi.org/10.1093/nar/gkz836, URL https://doi.org/10.1093/nar/gkz836, https://arxiv.org/abs/https://academic.oup.com/nar/article-pdf/48/D1/D941/31697918/gkz836.pdf

Fischl B (2012) Freesurfer. NeuroImage 62(2):774–781. https://doi.org/10.1016/j.neuroimage.2012.01.021

Fischl B, Sereno MI, Dale AM (1999) Cortical surface-based analysis. NeuroImage 9(2):195–207. https://doi.org/10.1006/nimg.1998.0396

Fortin JP, Cullen N, Sheline YI, et al (2018) Harmonization of cortical thickness measurements across scanners and sites. NeuroImage 167:104–120. https://doi.org/10.1016/j.neuroimage.2017.11.024

French L, Pavlidis P (2011) Relationships between gene expression and brain wiring in the adult rodent brain. PLoS Computational Biology 7(1). https://doi.org/10.1371/journal.pcbi.1001049

Froudist-Walsh S, Bliss DP, Ding X, et al (2021) A dopamine gradient controls access to distributed working memory in the large-scale monkey cortex. Neuron 109(21). https://doi.org/10.1016/j.neuron.2021.08.024

Garavan H, Bartsch H, Conway K, et al (2018) Recruiting the abcd sample: Design considerations and procedures. Developmental Cognitive Neuroscience 32:16–22. https://doi.org/10.1016/j.dcn.2018.04.004

Geschwind DH, Rakic P (2013) Cortical evolution: Judge the brain by its cover. Neuron 80(3):633–647. https://doi.org/https://doi.org/10.1016/j.neuron.2013.10.045, URL https://www.sciencedirect.com/science/article/pii/S0896627313009975

Glasser MF, Coalson TS, Robinson EC, et al (2016) A multi-modal parcellation of human cerebral cortex. Nature 536(7615):171–178. https://doi.org/10.1038/nature18933

Goulas A, Uylings HB, Hilgetag CC (2016) Principles of ipsilateral and contralateral cortico-cortical connectivity in the mouse. Brain Structure and Function 222(3):1281–1295. https://doi.org/10.1007/s00429-016-1277-y

Gómez-Robles A, Hopkins WD, Schapiro SJ, et al (2015) Relaxed genetic control of cortical organization in human brains compared with chimpanzees. Proceedings of the National Academy of Sciences 112(48):14,799–14,804. https://doi.org/10.1073/pnas.1512646112

Haak KV, Beckmann CF (2019) Plasticity versus stability across the human cortical visual connectome. Nature Communications 10(1). https://doi.org/10.1038/s41467-019-11113-z

Hagler DJ, Hatton S, Cornejo MD, et al (2019) Image processing and analysis methods for the adolescent brain cognitive development study. NeuroImage 202:116,091. https://doi.org/https://doi.org/10.1016/j.neuroimage.2019.116091, URL https://www.sciencedirect.com/science/article/pii/S1053811919306822

Hawrylycz M, Miller JA, Menon V, et al (2015) Canonical genetic signatures of the adult human brain. Nature Neuroscience 18(12):1832–1844. https://doi.org/10.1038/nn.4171

Hawrylycz MJ, Lein ES, Guillozet-Bongaarts AL, et al (2012) An anatomically comprehensive atlas of the adult human brain transcriptome. Nature 489(7416):391–399. https://doi.org/10.1038/nature11405

Homan P, Argyelan M, DeRosse P, et al (2019) Structural similarity networks predict clinical outcome in early-phase psychosis. Neuropsychopharmacology 44(5):915–922. https://doi.org/10.1038/s41386-019-0322-y

Horvát S, Gămănut, R, Ercsey-Ravasz M, et al (2016) Spatial embedding and wiring cost constrain the functional layout of the cortical network of rodents and primates. PLOS Biology 14(7). https://doi.org/10.1371/journal.pbio.1002512

Johnson WE, Li C (2007) Adjusting batch effects in microarray experiments with small sample size using empirical bayes methods. Batch Effects and Noise in Microarray Experiments p 113–129. https://doi.org/10.1002/9780470685983.ch10

Kong Xz, Liu Z, Huang L, et al (2015) Mapping individual brain networks using statistical similarity in regional morphology from mri. PLOS ONE 10(11). https://doi.org/10.1371/journal.pone.0141840

Kullback S, Leibler RA (1951) On information and sufficiency. The Annals of Mathematical Statistics 22(1):79–86. https://doi.org/10.1214/aoms/1177729694

Leming MJ, Baron-Cohen S, Suckling J (2021) Single-participant structural similarity matrices lead to greater accuracy in classification of participants than function in autism in mri. Molecular Autism 12(1). https://doi.org/10.1186/s13229-021-00439-5

Li J, Seidlitz J, Suckling J, et al (2021) Cortical structural differences in major depressive disorder correlate with cell type-specific transcriptional signatures. Nature Communications 12(1). https://doi.org/10.1038/s41467-021-21943-5

Li W, Yang C, Shi F, et al (2017) Construction of individual morphological brain networks with multiple morphometric features. Frontiers in Neuroanatomy 11. https://doi.org/10.3389/fnana.2017.00034

Markello RD, Arnatkevičiūtė A, Poline JB, et al (2021) Standardizing workflows in imaging transcriptomics with the abagen toolbox. eLife 10:e72,129. https://doi.org/10.7554/eLife.72129, URL https://doi.org/10.7554/eLife.72129

Markov NT, Ercsey-Ravasz MM, Ribeiro Gomes AR, et al (2012) A weighted and directed interareal connectivity matrix for macaque cerebral cortex. Cerebral Cortex 24(1):17–36. https://doi.org/10.1093/cercor/bhs270

Milham MP, Ai L, Koo B, et al (2018) An open resource for non-human primate imaging. Neuron 100(1):61–74.e2. https://doi.org/https://doi.org/10.1016/j.neuron.2018.08.039, URL https://www.sciencedirect.com/science/article/pii/S0896627318307682

Morgan SE, Seidlitz J, Whitaker KJ, et al (2019) Cortical patterning of abnormal morphometric similarity in psychosis is associated with brain expression of schizophrenia-related genes. Proceedings of the National Academy of Sciences 116(19):9604–9609. https://doi.org/10.1073/pnas.1820754116

Neale MC, Cardon LR (1994) Methodology for genetic studies of twins and families. Journal of Medical Genetics 67(9). https://doi.org/10.1136/jmg.30.9.800-a

Nielson DM, Pereira F, Zheng CY, et al (2018) Detecting and harmonizing scanner differences in the abcd study - annual release 1.0. bioRxiv https://doi.org/10.1101/309260, URL https://www.biorxiv.org/content/early/2018/05/02/309260, https://arxiv.org/abs/https://www.biorxiv.org/content/early/2018/05/02/309260.full.pdf

Perez-Cruz F (2008) Kullback-leibler divergence estimation of continuous distributions. In: 2008 IEEE International Symposium on Information Theory, pp 1666–1670, https://doi.org/10.1109/ISIT.2008.4595271

Romero-Garcia R, Atienza M, Clemmensen LH, et al (2012) Effects of network resolution on topological properties of human neocortex. NeuroImage 59(4):3522–3532. https://doi.org/10.1016/j.neuroimage.2011.10.086

Salvador R, Suckling J, Coleman MR, et al (2005) Neurophysiological architecture of functional magnetic resonance images of human brain. Cerebral Cortex 15(9):1332–1342. https://doi.org/10.1093/cercor/bhi016

Seidlitz J, Váša F, Shinn M, et al (2018) Morphometric similarity networks detect microscale cortical organization and predict inter-individual cognitive variation. Neuron 97(1). https://doi.org/10.1016/j.neuron.2017.11.039

Seidlitz J, Nadig A, Liu S, et al (2020) Transcriptomic and cellular decoding of regional brain vulnerability to neurogenetic disorders. Nature Communications 11(1). https://doi.org/10.1038/s41467-020-17051-5

Shen K, Bezgin G, Schirner M, et al (2019) A macaque connectome for large-scale network simulations in thevirtualbrain. Scientific Data 6(1). https://doi.org/10.1038/s41597-019-0129-z

Taquet M, Smith SM, Prohl AK, et al (2020) A structural brain network of genetic vulnerability to psychiatric illness. Molecular Psychiatry 26(6):2089–2100. https://doi.org/10.1038/s41380-020-0723-7

Vainik U, Paquola C, Wang X, et al (2020) Heritability of cortical morphology reflects a sensory-fugal plasticity gradient. bioRxiv https://doi.org/10.1101/2020.11.03.366419, URL https://www.biorxiv.org/content/early/2020/11/04/2020.11.03.366419, https://arxiv.org/abs/https://www.biorxiv.org/content/early/2020/11/04/2020.11.03.366419.full.pdf

Verhulst B, Prom-Wormley E, Keller M, et al (2018) Type i error rates and parameter bias in multivariate behavioral genetic models. Behavior Genetics 49(1):99–111. https://doi.org/10.1007/s10519-018-9942-y

Váša F, Seidlitz J, Romero-Garcia R, et al (2017) Adolescent tuning of association cortex in human structural brain networks. Cerebral Cortex 28(1):281–294. https://doi.org/10.1093/cercor/bhx249

Vértes PE, Rittman T, Whitaker KJ, et al (2016) Gene transcription profiles associated with inter-modular hubs and connection distance in human functional magnetic resonance imaging networks. Philosophical Transactions of the Royal Society B: Biological Sciences 371(1705). https://doi.org/10.1098/rstb.2015.0362

Wang H, Jin X, Zhang Y, et al (2016) Single-subject morphological brain networks: Connectivity mapping, topological characterization and test–retest reliability. Brain and Behavior 6(4). https://doi.org/10.1002/brb3.448

Wang Z, Scott DW (2019) Nonparametric density estimation for highdimensional data—algorithms and applications. WIREs Computational Statistics 11(4). https://doi.org/10.1002/wics.1461

Warrier V, Kwong AS, Luo M, et al (2021) Gene–environment correlations and causal effects of childhood maltreatment on physical and mental health: A genetically informed approach. The Lancet Psychiatry 8(5):373–386. https://doi.org/10.1016/s2215-0366(20)30569-1

Warrier V, Cliquet F, Grove J, et al (2022a) Genetic correlates and consequences of phenotypic heterogeneity in autism. Nature Genetics https://doi.org/10.1038/s41588-022-01072-5

Warrier V, Stauffer EM, Huang QQ, et al (2022b) The genetics of cortical organisation and development: A study of 2,347 neuroimaging phenotypes. arXiv https://doi.org/10.1101/2022.09.08.507084

Whitaker K, Vértes P, Romero-Garcia R, et al (2017) Adolescence is associated with genomically patterned consolidation of the hubs of the human brain connectome. Biological Psychiatry 81(10). https://doi.org/10.1016/j.biopsych.2017.02.390

Xu T, Nenning KH, Schwartz E, et al (2020) Cross-species functional alignment reveals evolutionary hierarchy within the connectome. NeuroImage 223:117,346. https://doi.org/10.1016/j.neuroimage.2020.117346, bALSA link https://balsa.wustl.edu/reference/976nz

Yang J, Lee SH, Goddard ME, et al (2011) Gcta: A tool for genome-wide complex trait analysis. The American Journal of Human Genetics 88(1):76–82. https://doi.org/10.1016/j.ajhg.2010.11.011

Zhang Y, Ma M, Xie Z, et al (2021) Bridging the gap between morphometric similarity mapping and gene transcription in alzheimer’s disease. Front Neurosci https://doi.org/10.21203/rs.3.rs-348434/v1

