## Supplementary Materials for "MIND Networks: Robust Estimation of Structural Similarity from Brain MRI"

##### *Overview*

In this supplement, we provide additional information to illustrate our workflow, more comprehensively characterize MIND networks and MSNs, and report sensitivity analyses and other results aimed at contextualizing the main text as follows:

- Figures S1 and S2: additional figures to clearly illustrate the pipeline for constructing a MIND network from a raw T1w image, and to visualize our subject inclusion criteria at each stage of analysis.
- Figures S3, S4, and S5: additional analyses and figures characterizing MIND networks and MSNs, including their connection to Euclidean distance and their association with brain size.
- Figures S6 and S7: additional sensitivity analyses concerning the coupling between networks of transcriptional similarity from the Allen Human Brain Atlas (AHBA) and both MIND networks and MSNs.
- Figure S8 and section "Replicating existing heritability results": additional results relating to MIND network heritability to better contextualize our results in the context of published work, and to describe how the patterning of MIND heritability corresponds to neuroanatomical features and MIND network phenotypes.

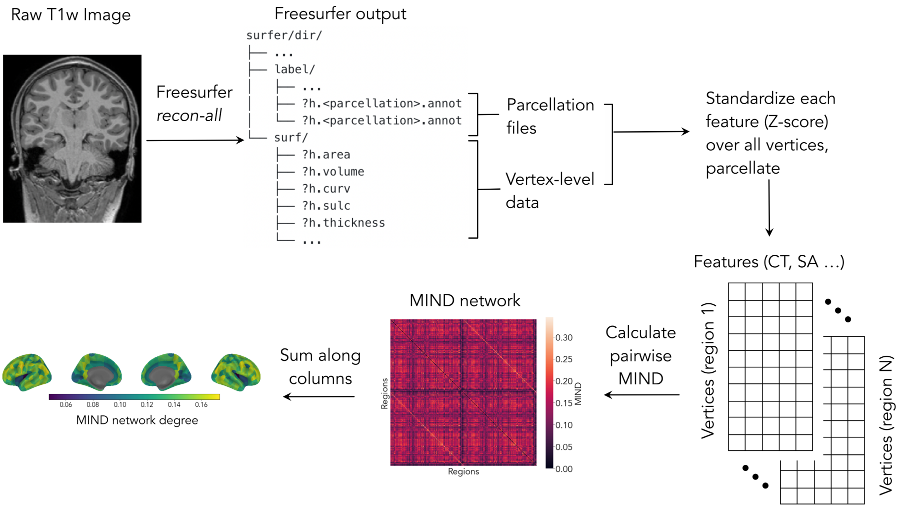


**Fig. S1** A visualization of the entire MIND pipeline, from raw T1w image to MIND network phenotypes, using FreeSurfer as the image-processing tool of choice.

### Workflow

#### *MIND, from image to phenotype*

Figure S1 demonstrates each step of the workflow from the raw data of a T1w image into final MIND network phenotypes, using FreeSurfer as the image processing tool of choice. Using the T1w nifti file as input, we process the scan using FreeSurfer's `recon-all` command. This outputs `label/` and `surf/`, the folders in the surfer directory containing all necessary information for MIND calculation. The `surf/` folder houses all vertex-level data, with `?h.area`, `?h.volume`, `?h.curv`, `?h.sulc`, and `?h.thickness` containing the vertex-level values of surface area, grey matter volume, mean curvature, sulcal depth, and cortical thickness, respectively. The `surf/` folder houses the `?h.annot` files necessary to parcellation the vertex-level data. If the use of a non-standard parcellation is required, additional `?h.annot` files will need to be created using e.g. the `mris_ca_label` command. Given these outputs, we standardize (Z-score) each feature across all vertices, then parcellate the vertex data to create data-frames of vertices of all features per region. Calculating the MIND similarity statistic then produces the final MIND network, and the weighted nodal degree is then computed as the sum or average along the columns.

#### *Data Inclusion*

We provide a visualization of the exact breakdown of the subjects used at each stage of our analysis in Figure S2. As shown, some subjects were extreme outliers in terms of data quality, measured by Euler index. To remove such scans, we set a cutoff threshold of -120 as shown, which corresponded to a median absolute deviation (MAD) score of 2.6.

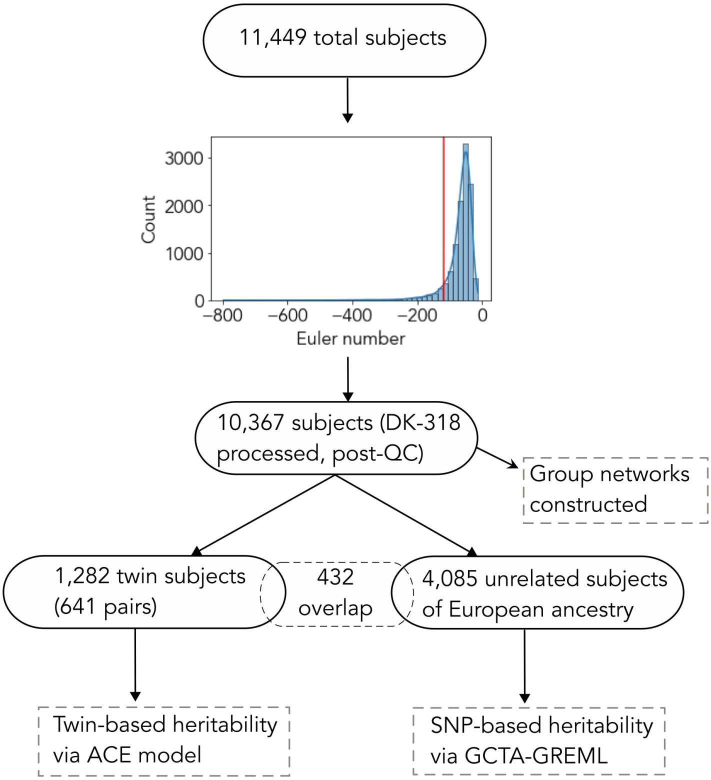


**Fig. S2 Subjects included for each analysis.** Flow chart indicating the total number of subjects included for construction of the group-level networks, and for both heritability analyses. The histogram shows the Euler number QC cutoff threshold of -120. The 10,367 post-QC subjects listed were those who had a) an Euler number over -120 and b) no regions assigned zero vertices in the DK-318 parcellation.

#### Characterizing MIND networks and MSNs

##### *Relationship to Euclidean distance*

Building on previous work demonstrating an exponentially decaying relationship between Euclidean distance and both structural (Goulas et al, 2016; Horvát et al, 2016) and functional (Salvador et al, 2005) brain network connections, we studied the relationship between distance and structural similarity for both MIND networks and MSNs. Replicating the results from Seidlitz et al (2018), we found a exponentially decaying relationship for MSNs, and additionally observed a similar relationship between distance and MIND network edges as shown in Figure S3.

##### *Consistency across cortical parcellations*

In Figure S4, we provide a more detailed visualization of the result that MIND networks demonstrate much higher consistency across cortical parcellations

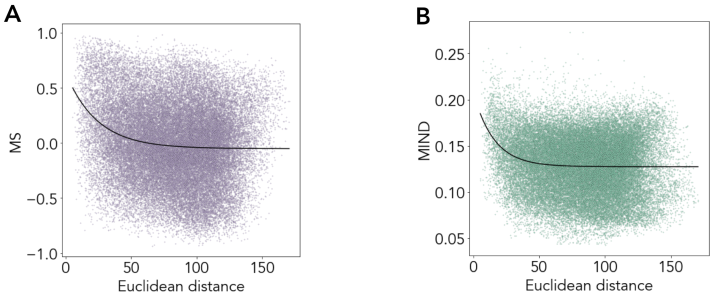


**Fig. S3 Relationship to Euclidean distance.** The exponentially decaying relationship between Euclidean distance and edge strength for group-level morphometric similarity networks (A) and MIND networks (B).

compared to MSNs, discussed and reported in Figure 2H. In this figure, we additionally show the cortical maps of network weighted degree for MIND networks and MSNs and the weighted degree distribution for each method and parcellation, shown on the diagonal of the pairplots in Figure S4 B and D.

##### *Relationship to Total Intracranial Volume*

Given that brain size is the most salient axis of variation in brain structure, we expected that measures of structural similarity should demonstrate a significant relationship with eTIV. We found that MIND networks and MSNs have different relationships to brain size, as measured by estimated total intracranial volume (eTIV); specifically MIND network features more closely reflect variation in brain size at the individual level (Figure S5). At the degree level, MIND network degree is on average positively correlated with total eTIV (mean $r = 0.23$), though the extent of this correlation varies dramatically across the cortex (Figure S5A, C). By contrast, MSN degrees are also differentially related to volume across the cortex, though the distribution is centered on zero (Figure S5B, D). Finally, we observe a strong difference in the relationship of total eTIV with global MIND vs. global MS (where global network measures are simply calculated as the sum of all entries in the similarity matrix). These global values are strongly related to eTIV in MIND networks ($r = 0.59$), but not in MSNs ($r = 0.04$) (Figure S5E, F). In future work, these relationships will be important to consider when interpreting results – for example, by ensuring that regional differences in MIND observed in a case-control study are not due exclusively to differences in brain volume.

#### AHBA additional analysis

##### *Sensitivity analysis over genes and donors used*

We assessed the stability of the high correlation between the group-level MIND and transcriptomic similarity networks by performing extensive sensitivity analysis. We first recreated the results over a range of gene-filtering steps, using an increasingly restrictive threshold of differential stability as the gene

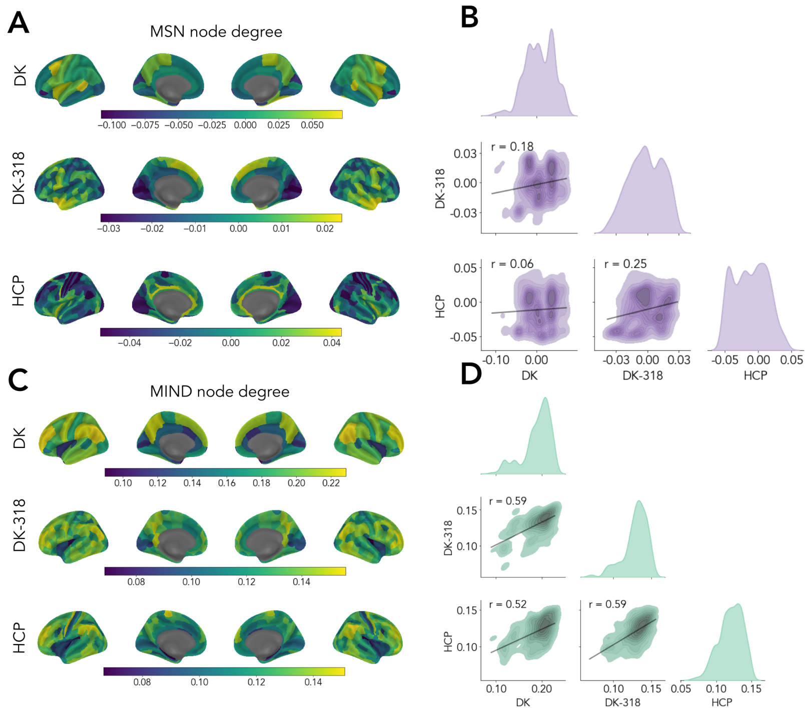


**Fig. S4 Robustness to changes in parcellation.** A, C) The weighted node degree of MIND networks and MSNs for Desikan Killiany (DK), DK-318, and HCP parcellations. B, D) Association between weighted node degrees across all pairs of DK, DK-318, and HCP parcellations. The correlations were calculated by mapping from regional to vertex space as described in the main text. These plots present a more detailed visualization of the same values reported in Figure 2F. Diagonal entries show the degree distribution for each network type and parcellation.

inclusion criterion (Hawrylycz et al, 2015). Differential stability measures the mean pairwise correlation of the expression patterns between donors; higher thresholds therefore indicate more conserved spatial patterning. As shown in Fig. S6A, the correlation between structural similarity and transcriptomic similarity was highly conserved across the full set of filtering steps, even in the most restrictive case which included only the most stable 10% of genes.

Next, we studied the sensitivity of the results from Fig. 4A-C to the number and identity of the donors included in the gene similarity construction. This analysis was introduced in the main text (Fig. 4D), and is extended here. Using the set of 29 regions covered by all 6 donors, we constructed transcriptomic similarity networks for all possible donor subsets. The pattern of correspondence was highly stable, with a slight and monotonic gain as more donors were included, reflecting a gene-structure connection that becomes stronger as the signal to noise ratio improves. Interestingly, even when considering gene similarity networks generated from each donor independently, the variance of the correlation with MIND was relatively low. Thus, despite the small sample size

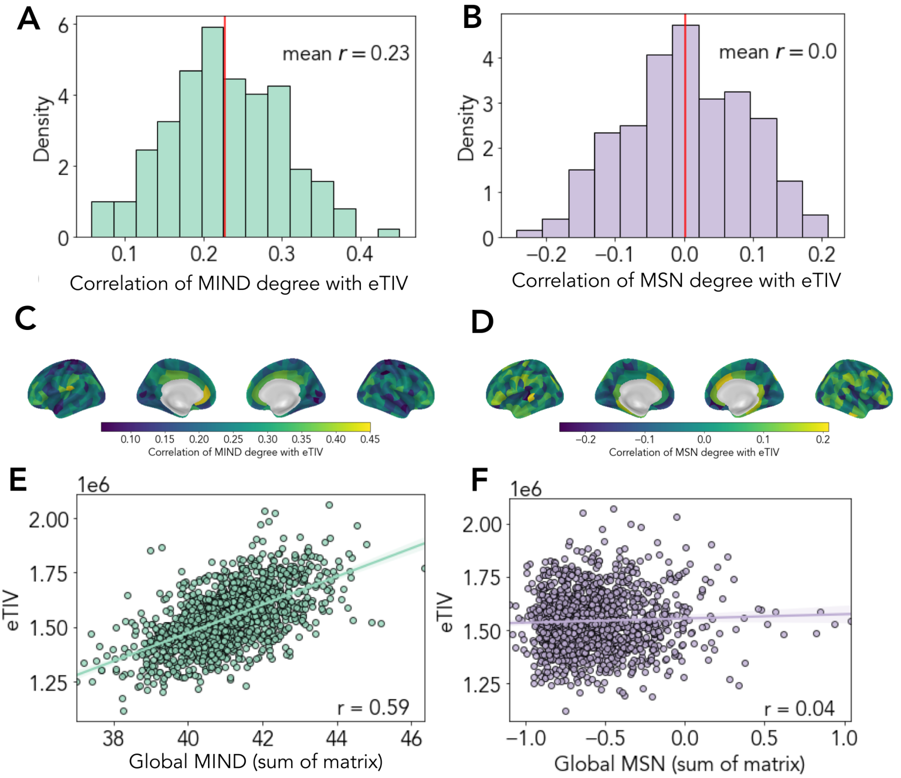


**Fig. S5 Relationship between MIND, MS, and eTIV.** A-B) The distribution of correlations of regional node degree for MIND networks and MSNs with total GMV. C-D) A visualization of the spatial organization of the correlations reported in C and D on the cortex. E-F) the correlation between eTIV and global MIND and MS. Global network measures were calculated as the sum over the entire similarity matrix.

from the AHBA, these results point to a shared and replicable structural and transcriptomic cortical architecture in the general population.

##### *Replication with DK-318*

We repeated the main analysis comparing structural and transcriptional similarity using the DK-318 atlas, which is closely related to the DK-308 atlas to compare AHBA gene expression with changes in MSNs under various diesease (Morgan et al, 2019; Seidlitz et al, 2020; Li et al, 2021; Zhang et al, 2021). Because each region in this parcellation is smaller, few regions are covered by all 6 donors, with many only covered by 1-2 donors. Processing steps, such as gene or regional filtering, thereby influence the results more heavily than when using the coarser DK atlas, which has higher donor coverage within each region. In addition to filtering for genes based on differential stability, we therefore additionally considered filtering regions based on the number of donors with gene probe coverage within it. The results over a full grid-search

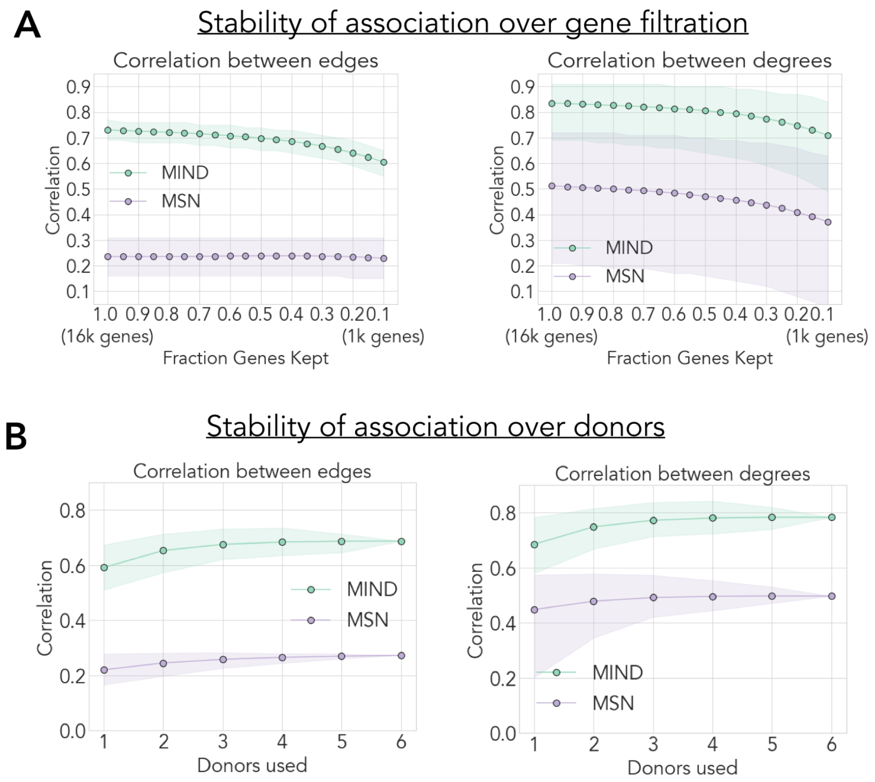


**Fig. S6 Stability of association of structural similarity and gene coexpression networks**. A) Stability of the correlation between structural and transcriptomic networks over genes included. Gene networks were constructed using genes filtered based on differential stability (DS) at a range of thresholds, from 100% to 10% inclusion, in 5% increments. Shading represents 95% CI. B) Stability of the correlation between structural and transcriptomic networks over AHBA donors included. The first plot is replicated from Figure 4D for ease of comparison. For each number of donors included, all combinations of gene similarity networks were constructed (no gene filtering) and the mean edge and degree correlation were calculated. There were $\binom{6}{n}$ possible networks created for $n$ included donors. Shading indicates the minimum and maximum value of the association observed at each number of included donors.

sweep over combinations of gene- and region-filtering steps are shown in Fig. S7. Overall, at both the degree and edge level, the original result is reproduced that MIND networks demonstrate much higher correspondence with the gene similarity networks.

#### Additional heritability analyses

##### *Replicating existing heritability results*

In previous work on the heritability of GMV in the ABCD dataset by Bethlehem et al (2022), heritability estimates were reported using the path

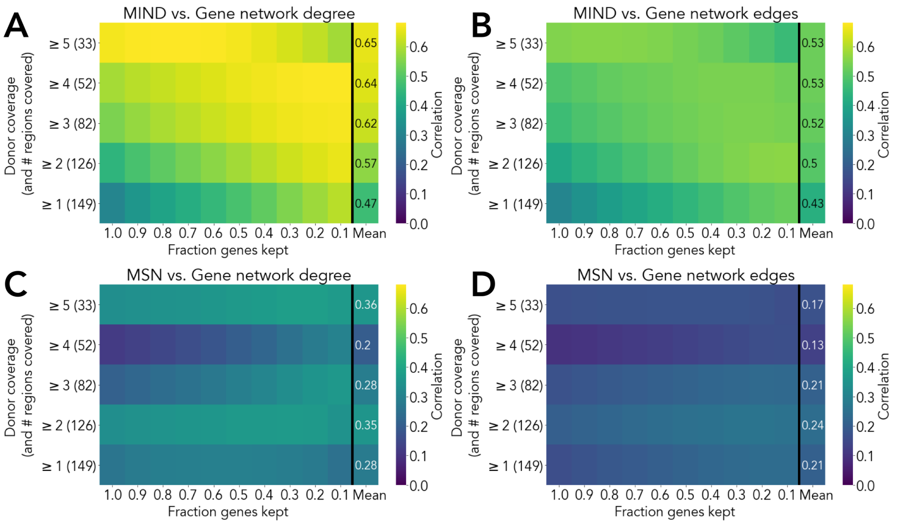


**Fig. S7 Correspondence between structural and transcriptomic similarity networks, DK-318**. Heatmaps showing structural vs. transcriptomic degree (A, C) and edge (B, D) correlations using the DK-318 parcellation. Gene networks were constructed over a grid of gene filtering (100%- 10% inclusion using DS) and regional filtering ($\geq$1-donor coverage to $\geq$5-donor coverage) steps. Structural networks were associated at each combination of filtration steps. The rightmost column of each heatmap reports the mean association at each level of regional filtering. As only 6 regions (3.8%) were covered by all 6 donors, we did not consider filtering only for regions with complete 6-donor coverage.

coefficients $a, c$ and $e$ from structural equation modeling, rather than the interpretation of heritability as the proportion of variance reported in the main text, calculated as $\frac{a^2}{a^2+c^2+e^2}$ (Verhulst et al, 2018). Therefore to compare our results to these previously published path coefficients, we estimated the path coefficients for total brain GMV, comparing them to those reported by Bethlehem et al (2022). The published coefficients were the following: $a = 0.745, c = 0.624, e = 0.238$, and we calculated values of $a = 0.765, c = 0.596, e = 0.246$, which is highly consistent.

##### *Contextualizing heritability patterns*

Figure S8 provides additional material to contextualize the heritability results from the main text. Having calculated $h^2_{twin}$ for all regional metrics and weighted network degree for both structural similarity network types, we found that the patterns of heritability were positively correlated for all pairs of regional features (Fig. S8A). Just as we found a negative correlation between MIND edge strength and $h^2_{twin}$, we observed a negative relationship between MIND weighted nodal degree and $h^2_{twin}$ (Fig. S8B), with differences observed across von Economo classes (Fig. S8C). Figure S8D shows the distribution of MIND network degree by different cytoarchitectonic (von Economo) classes,

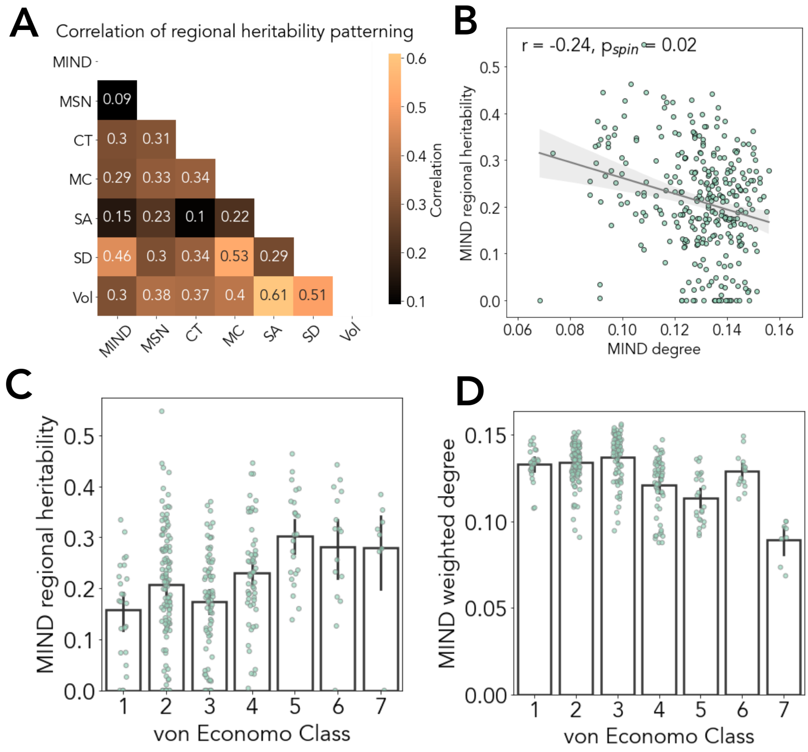


**Fig. S8** A) Correlation matrix of the relationship between heritability patterns across the cortex for all studied regional features, showing a positive correlation between all vectors of regional heritabilities. B) Scatterplot of $h^2_{twin}$ versus MIND network weighted degree, with fitted line indicating significant negative correlation; each point is node in the network. C) Twin-based heritability of MIND network weighted degree by different cytoarchitectonic (von Economo) classes: 1, agranular cortex, primary motor cortex; 2, association cortex; 3, association cortex; 4, secondary sensory cortex; 5, primary sensory cortex; 6, limbic regions; 7, insular cortex (Seidlitz et al, 2018). D) MIND network degree by different cytoarchitectonic (von Economo) classes.

showing the insular and primary sensory cortices to have the lowest average MIND weighted degree in contrast to the patterning of MIND degree heritabilities.
